# Intrarenal myeloid subsets associated with kidney injury are comparable in mice and patients with lupus nephritis

**DOI:** 10.1101/2023.06.24.546409

**Authors:** Paul J. Hoover, David J. Lieb, Joyce Kang, Stephen Li, Michael Peters, Chirag Raparia, Arnon Arazi, Thomas Eisenhaure, Saisram S. Gurajala, Qian Xiao, Rakesh Mishra, Max Spurrell, Rajasree Menon, Matthias Kretzler, Jonathan Chen, Linda Nieman, Abraham Sonny, Dawit Demeke, Jeffrey Hodgin, Joel Guthridge, Andrea Fava, Robert Clancy, Chaim Putterman, Peter Izmirly, H. Michael Belmont, Kenneth Kalunian, Diane Kamen, David Wofsy, Jill Buyon, Judith A. James, Michelle Petri, Betty Diamond, Soumya Raychaudhuri, The Kidney Precision Medicine Project, The Accelerating Medicines Partnership: RA/SLE network, Nir Hacohen, Anne Davidson, Co-senior

## Abstract

Resident macrophages and infiltrating monocytes in kidneys of patients with lupus nephritis are altered both in frequency and function relative to their counterparts in healthy kidneys. The extent to which mouse models might be useful in developing approaches to target these cells for treating lupus nephritis is poorly understood. Here, we studied four common lupus mouse models that share clinical, serologic, and histopathologic kidney changes with humans. Using single-cell profiling and multiplex spatial imaging to analyze the intrarenal myeloid compartment with the onset of clinical disease in these models, we identified monocyte and macrophage subsets that expand or contract in kidneys with clinical nephritis. A unique subset of classical monocytes expanded with the onset of disease and expressed genes such as *CD9, Spp1, Ctsd, Cd63, Apoe,* and *Trem2* that were previously shown to be induced by tissue injury and play a role in inflammation, lipid metabolism and tissue repair in other organs. Resident macrophages transitioned from a pro-inflammatory to a similar injury-associated state with onset of disease. To test whether these findings in mouse models were also observed in humans, we re-analyzed monocytes and macrophages in a single-cell RNAseq dataset of kidney biopsies from 155 patients with lupus nephritis and 30 healthy donors, collected by the NIH AMP RA/SLE consortium. Human monocytes and macrophages showed conserved changes in gene expression programs associated with lupus nephritis disease indices, and localized to similar kidney microenvironments as in mice. By identifying myeloid subsets and disease-associated alterations in biological processes that are conserved across species, we provide a strong rationale for functional studies of these cells and pathways in mice to uncover mechanisms and find targets relevant to human lupus nephritis.

**One sentence summary:** This study characterizes intrarenal myeloid cells from four lupus mouse models and 155 patients with lupus nephritis using single-cell RNA-seq and imaging, and identifies novel infiltrating and resident myeloid subsets that are conserved between mouse and human lupus nephritis, thus providing a map and strong rationale for functional studies in mice with relevance to human disease.

## INTRODUCTION

Systemic lupus erythematosus (SLE) affects >200,000 Americans, of whom >40% will develop lupus nephritis and 8-10% will develop renal failure over 20 years (*1*) (*2*) (*3*). SLE is caused by the loss of immune tolerance and the production of autoantibodies against nuclear material. Intrarenal deposition of IgG and complement triggers kidney inflammation and pathologic changes in the glomerulus and tubulointerstitium, resulting in functional tissue injury (*4*) (*5*).

Myeloid cell infiltration has been associated with kidney injury and fibrosis in patients with lupus nephritis (*6*). Using single-cell RNA sequencing (scRNA-seq), we previously reported five novel myeloid subsets from the kidney biopsies of 23 unique lupus nephritis patients and 10 healthy donors (*7*). More recently, scRNA-seq analyses of kidney biopsies from 155 unique lupus nephritis patients (*3*) captures the most comprehensive view of this immune compartment to date and will be described in an upcoming resource study. The role of these immune cells has been difficult to study due to the inability to perform functional studies on small numbers of cells isolated from human kidney biopsies.

Mouse models of lupus have been used to investigate the molecular and cellular mechanisms that underpin lupus and lupus nephritis, and harbor multiple myeloid cell types in the kidney (*6*). Like humans, lupus mouse models are characterized by antinuclear autoantibodies and immune complex-mediated glomerulonephritis with variable histopathological changes in the glomerulus and tubulointerstitium that result in progression to irreversible kidney injury (*4*) (*8*). To better utilize these mice, we need to pinpoint which molecular and cellular features of the models reflect human disease.

In this study, we aimed to characterize intrarenal myeloid cells from four lupus mouse models and to determine if there are comparable patient myeloid cell states, which will enable using lupus mouse models to study cell types and pathways most relevant to humans. The four mouse models have distinct pathogenic mechanisms and anti-nuclear antibody subtypes that reflect different facets of human lupus nephritis (**Fig. 1A**) (*9*). Two models overexpress the *Tlr7* (*Yaa* locus in males) and develop early and severe (NZW/BXSB) or later and more indolent onset (Sle1.Yaa) lupus nephritis. The additional copy of *Tlr7* enhances the production of anti-chromatin and anti-RNA antibodies and is critical for driving murine lupus nephritis in the background of other predisposing alleles in these strains (*10*) (*11*). In humans, single nucleotide polymorphisms (SNPs) linked to *TLR7* increase the risk of SLE and lupus nephritis (*12*), and TLR7 gain of function causes lupus (*13*). We used two additional lupus nephritis strains that are characterized by the expansion of lymphoid cells in the setting of normal *Tlr7* expression. MRL/lpr mice are a spontaneous model in which *Fas* deletion in the permissive MRL genetic background causes both B cell and DN T cell expansion and production of multiple autoantibodies that drive severe proliferative and interstitial lupus nephritis. Kidney-infiltrating double negative T cells and T cells producing IFNγ or IL-17 are present in this model (*14*) (*15*) and are also found in human lupus nephritis (*7*). NZB/W mice produce anti-dsDNA antibodies and harbor genetic loci that predispose to SLE and that are shared with SLE patients (*16*) (*17*) (*18*).

**Figure 1.**
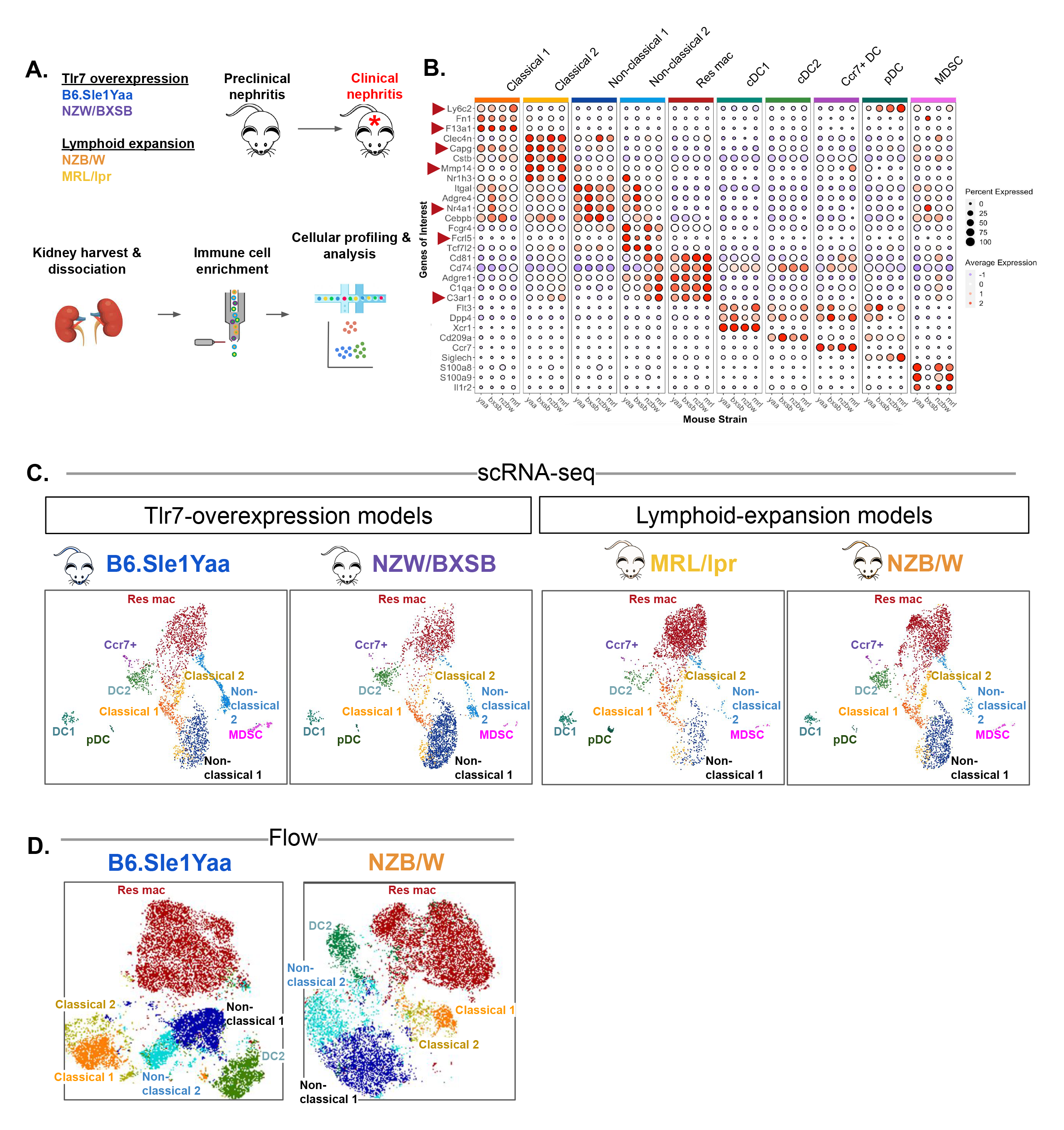
Summary of the intrarenal myeloid states identified in 4 lupus mouse strains. **(A**) Schematic of experimental workflow. Single cell suspensions were prepared from kidneys harvested from 4 lupus mouse strains with preclinical or clinical nephritis for analysis by single cell transcriptomic profiling or flow cytometry. Clinical nephritis was defined by fixed proteinuria of >300 mg/dl for >2 weeks and was confirmed by light microscopy. **(B)** Dot plot showing the expression of enriched cell-type canonical markers used to identify 10 intrarenal myeloid subsets. Genes used for RNA FISH staining (Sup. Figs 8, 9, 22, 23) are marked. **(C**) Individual UMAP plots of intrarenal myeloid cells from the integrated analysis of preclinical and clinical nephritis for each strain (n=2,993 cells in each UMAP). (**D**) tSNE plots of flow cytometry depicting intrarenal myeloid subsets from preclinical or clinical nephritis in B6.Sle1.Yaa. Each plot contains 24,000 cells from the CD45+ CD11b+ XCR1- Ly6G- gate (6,000 cells from each mouse).

We compared the molecular and spatial landscape of the myeloid compartment in the kidneys of these four mouse models of lupus nephritis with those of renal biopsy samples from 155 lupus nephritis patients and 30 healthy controls. By aligning intrarenal myeloid cell transcriptional profiles across mouse and human lupus, we mapped similar myeloid subsets between species and identified comparable myeloid states associating with disease, including multiple subtypes of monocytes, macrophages, and dendritic cells. These findings provide a roadmap for functional studies of specific mouse cell subsets that are mirrored in human lupus nephritis.

## RESULTS

### Intrarenal myeloid subsets identified by scRNA-seq and flow cytometry in four lupus mouse models

We used magnetic beads and FACS to enrich for viable total hematopoietic (*CD45*+) or myeloid (*CD11b*+ or *CD11c*+) cells from dissociated kidneys and blood of four lupus mouse models before (‘preclinical nephritis’ defined by proteinuria ≤30 mg/dl) or after (‘clinical nephritis’) the development of clinical disease defined by kidney dysfunction (proteinuria of ≥300 mg/dl for >2 weeks) (*19*) (**Fig. 1A**). We used the 10x scRNA-seq platform to measure single-cell transcriptomes and applied standard bioinformatic approaches to identify high quality cells. We focused downstream analysis on *Cd45+* clusters with expression of myeloid (*Cd68*, *Csf1r*, or *Flt3*) but not T and B (*Cd3e*, *Ms4a1*) markers (**Fig. S1A**). Using Louvain clustering, we partitioned the resulting 20,299 kidney myeloid cells into 10 subtypes of monocytes, macrophages, and dendritic cells across four models (**Fig. 1B, C, S1B**). Using canonical and subset-specific myeloid markers from our scRNA-seq analysis, we created a discriminative 16-color flow cytometry panel to further quantify the proportions of the most frequent myeloid subsets from kidneys and blood in independent Sle1.Yaa and NZB/W mice (**Fig. 1D, S2**). Below we primarily focus our analyses on changes in monocytes and resident macrophages that are associated with disease progression.

### An injury-associated classical monocyte subset is expanded in mice with clinical nephritis

We identified two subtypes of classical monocytes, C1 and C2. C1 monocytes (Ly6C^hi^/Ccr2^hi^) were found in blood and kidneys of preclinical and clinical nephritic mice in all 4 mouse models, while C2 monocytes were rare in the blood and expanded in kidneys of Sle1.Yaa and NZB/W mice with clinical disease, based on flow cytometry (p<0.01) (**Fig. 2A,C, S3**) and consistent with scRNA-seq (p<0.01) (**Fig. 2B, D, E, S4A**). This population could be distinguished from classical 1 monocytes in flow cytometry by expression of *Clec4n* and lower expression of *Ly6c2* and *Ccr2* (**Fig. S2A, B**). C1 and C2 were highly related transcriptionally based on partition-based graph abstraction (PAGA) (**Fig. 2F**) (that has been used to reconstruct lineage relationships among cell states) (*20*). Thus, a simple hypothesis is that C1 infiltrates from the blood and differentiates from C1 to C2 in the kidney. Relative to C1, C2 monocytes were enriched in genes involved in uptake and degradation of extracellular molecules, lipid metabolism and complement (**Fig. 2G**) and expressed known markers of injury-induced monocytes *CD9, Spp1, Ctsd, Cd63, Apoe, Fth1, Ftl1,* and *Fabp5* (**Fig. 2H**) (as well as *Gpnmb* and *Trem2* in a subset of C2, (**Fig. S4B**), which participate in tissue repair and regulation of fibrosis and inflammation in multiple organs (*21*) (*22*) (*23*). Thus, C2 monocytes correspond to known injury-associated cells and expand in mice with lupus nephritis.

**Figure 2.**
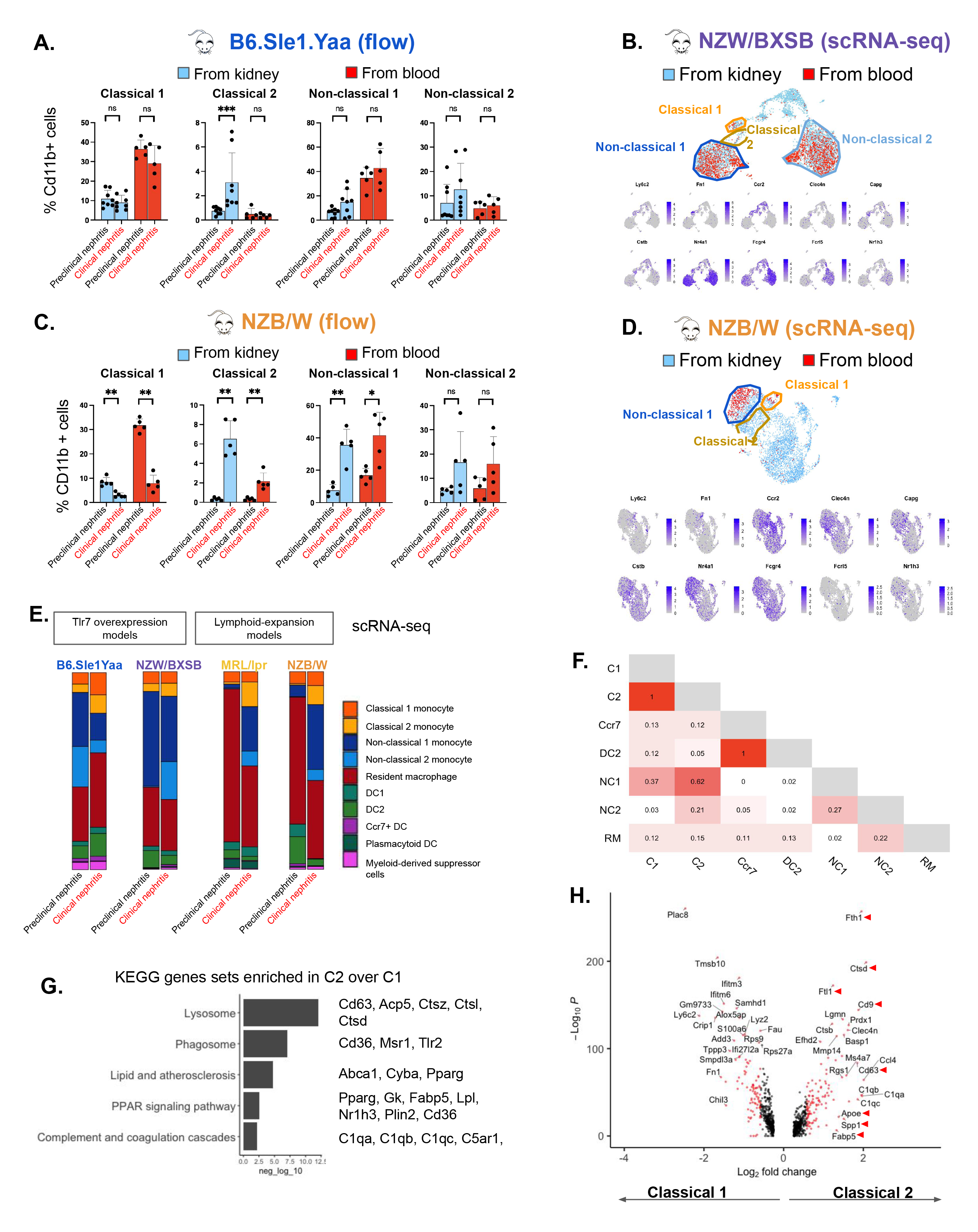
Summary of classical and non-classical monocytes identified in 4 lupus mouse strains. **(A) & (C)** Percentage of Cd11b+ cells identified in flow cytometry as Classical 1, Classical 2, Non-classical 1, Non-classical 2 in preclinical or clinical nephritis. Cells were collected from CD45 enriched kidney cell suspensions (light blue) and matched blood (red) from B6.Sle1.Yaa (A) and NZB/W (C) mice. 5 - 8 mice per condition. (ns: p > 0.05; *: p < 0.05; **: p < 0.01; ***: p < 0.001; ****: p < 0.0001). (**B) & (D)**. Top panel: Co-clustering of single myeloid cells collected from CD45 enriched kidney cell suspensions (light blue) and matched blood (red) from NZW/BXSB (B) and NZB/W (D) mice during clinical nephritis. Bottom panels in (**B**) & (**D**) feature plots of canonical markers for identifying Classical 1, Classical 2, Non-classical 1, Non-classical 2 monocytes. (**E)** Normalized bar plots comparing myeloid subset frequencies in preclinical or clinical nephritis for each lupus mouse strain determined by scRNA-seq droplet proportions. (**F**) Pairwise comparison of PAGA transcriptional relatedness of myeloid subclusters subclusters. (**G**) KEGG gene sets enriched in C2 over C1 from mice. (**H)** Volcano plot depicting differentially expressed genes between C1 (left) and C2 (right). Labeling reflects differentially expressed genes identified in injury associated macrophages from other studies (see text).

### Non-classical monocytes split into two subtypes and associate with disease

Two clusters, NC1 and NC2, were annotated as non-classical monocytes based on low *Ly6c2*, *Ccr2* and high *Nr4a1*, *Cebpb*, and *Itgal* expression (*24*) (*25*) (**Fig. 1B, S1A, B**). Both clusters expanded in the blood (**Fig. 2A, C**) and kidneys (**Fig. 2A, C, E, S3A, B**) of mice with clinical lupus nephritis (with statistical significance for NZB/W and a similar trend for Sle1.Yaa). NC1 was strongly related to C2 and C1 by PAGA (**Fig. S2F**) consistent with the known development of non-classical from classical monocytes (*26*), while NC2 was equally related to NC1, C2 and resident macrophages, making its origin unclear. A cell type similar to NC2 was recently described in the spleens of Sle1.Yaa mice (*27*) but its role is not known. Comparing these subsets, NC1 cells were enriched for gene sets associated with phagocytosis and C-type lectin receptor signaling (**Fig. S5A**), while NC2 showed higher expression of programs for cell-adhesion and TGF-beta signaling (**Fig. S5A**) Fc receptors (*Fcgr4*, *Fcrl5*) and *Nr1h3* (**Fig. S5B**), a transcription factor that suppresses inflammatory genes and enhances clearance of ingested apoptotic cells (*28*) (*29*).

### Intrarenal resident macrophage states shift with the onset of disease

One major cluster, RM, was designated as resident macrophages because no cells resembled those from blood in this cluster, and it was not related to classical and non-classical monocytes based on PAGA (**Fig. 3F**). RM expressed known markers of resident macrophages including *Adgre1* (that encodes F4/80), *Cd81*, *Cd74 and C1qa* (*30*) (**Fig. 1B, S1B**). This population was among the most frequent before and after kidney injury in the 4 lupus models (**Fig. 3E, S3A, B**). Compared to monocytes, resident macrophages were enriched for MHC class II antigen presentation and clearance of cellular debris via complement (*C1qa*, *C1qb*, *C1qc)* and Fc gamma receptor-mediated phagocytosis (*Fcgr1*, *Fcgr2b*, *Fcgr3*, *Axl)* (**Fig. 1B, S1B**). RM split into 6 subclusters/states, with RM0 increasing (p<0.05) and RM1 decreasing (p<0.05) with disease (**Fig. 3A, S6A, B**). There was high transcriptional similarity between RM0 and RM1 (**Fig. 3C**), consistent with the possibility that they are related by differentiation. RM1 expressed chemokines (*Ccl3, Ccl4, Cxcl2*) and an anti-fibrotic gene, *Mmp13* (*31*) (**Fig. 3B, D)**. In contrast, RM0 expressed *Ctsd, Cd63, Apoe, Trem2, Ctsa, Ctsb, Acp5, Lgmn,* and *Pltp* (**Fig. 3B, D**), similar to classical 2 monocytes (above) and repair macrophages across different tissues and species (*22*) (*32*) (*33*) (*23*). Other RM clusters were less abundant than RM0 and RM1 (Fig. 3A), and included: RM2 that expressed genes known to induce RAGE-dependent apoptosis (*S100a6*) (*34*), promote phagocytosis (*Sirpb1c*) (*35*), inhibit kidney monocyte infiltration (*Ccl6*) and fibrosis (*Ccl9*) (*36*) (**Fig. 2B**); RM3 expressed interferon-stimulated genes and chemokines and the Cd40 receptor. RM4 was infrequent and differentially expressed *Hpgd* that is known to metabolize pro-PGE2 to an anti-inflammatory molecule (*37*), *Stab1* that enables tissue repair by limiting fibrosis (*38*), and *Gas6* that is anti-inflammatory during acute inflammation and pro-fibrotic in chronic conditions (*39*). RM5 was also infrequent and was enriched for *Spp1* that marks pro-fibrotic macrophages (*40*) (**Fig. 3B**). We note that a small number of proliferating resident macrophages were present before and after clinical nephritis that could contribute to resident macrophage shifts (**Fig. S6C**). To summarize, we find that kidneys from mice with lupus nephritis are dominated by RM0 macrophages that express an injury-associated signature associated with tissue repair.

**Figure 3.**
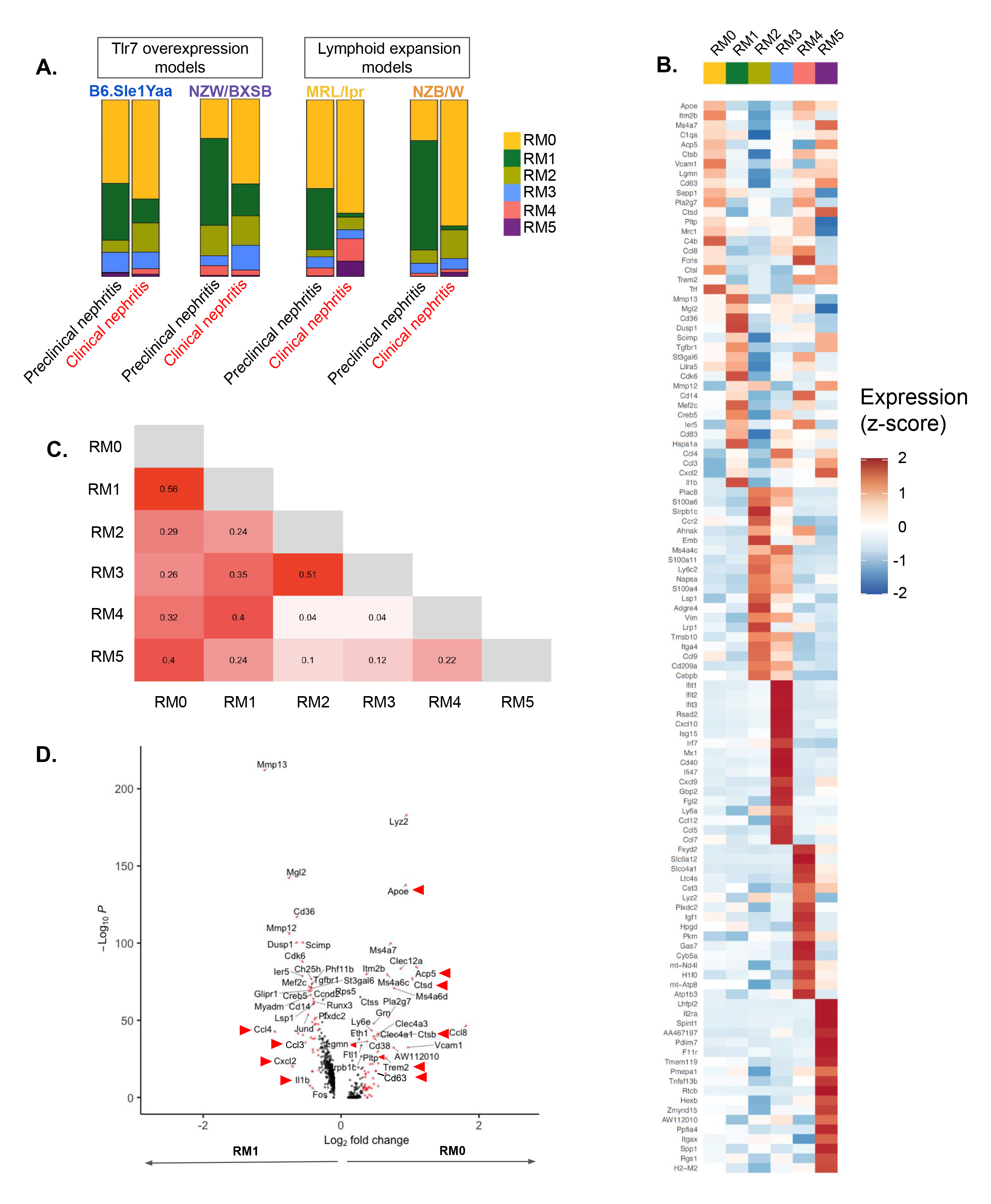
Summary of the intrarenal resident macrophage subclusters identified in 4 lupus mouse strains. **(A)** Normalized bar plots comparing frequencies of resident macrophage subclusters from each strain in preclinical or clinical nephritis measured by scRNA-seq droplet proportions. (**B**) Heatmap showing the scaled expression of top 20 discriminative genes for each resident macrophage subcluster. Color scheme is based on z-score, from −2 (blue) to 2 (red). (**C**) Pairwise comparison of PAGA transcriptional relatedness of resident macrophage subclusters. (**D**) Volcano plot depicting differentially expressed genes between RM1 (left) and RM0 (right). Labeling reflects differentially expressed genes identified in injury-associated macrophages from other studies (see text).

### *In-situ* localization of monocytes and macrophages in mouse kidneys

To understand where in the kidney monocyte and macrophages may perform their functions, we mapped their positions in Sle1.Yaa kidney sections collected before and after clinical disease. We used RNA FISH probes targeting *Csf1r* transcripts in order to identify monocytes and macrophages and 2-3 probes targeting cluster-specific genes based on our scRNA-seq analysis (**Fig. 1B**) to localize subsets (**Fig. S7**), noting that no individual gene perfectly marked a particular cluster. C1 monocytes (marked by *Csf1r, Ly6c2, F13a1*) were infrequent but localized to the glomerulus (**Fig. S8A**), near vessels and immune aggregates in clinical nephritis (**Fig. S9A**). C2 monocytes (*Csf1r, Capg, Mmp14)* localized to glomerular, peri-glomerular sites (**Fig. S8B**) and within immune infiltrates (**Fig. S9B**) around blood vessels and in the tubulointerstitial space. Both NC1 and NC2 localized to glomerular and periglomerular sites (**Fig. S8C**), were rarely found in dense infiltrates, and were rare in the tubulointerstitium and medulla (**Fig. S9C**). Resident renal macrophages (*Csf1r, C3ar1)* expanded in periglomerular regions and in the medulla in clinical disease (**Fig. S8D, S9D**), consistent with our and prior work (*41*) (*42*) (*43*). We note that some of the RM staining in glomerulus (**Fig. S8D**) may also label C2 cells since they express lower levels of *C3ar1* (**Fig. 1B**). We conclude that C1 and NC monocytes are restricted to the glomeruli whereas cells expressing injury-associated programs are located within the nephritic glomerulus (C2 monocytes) and tubulointerstitial space (C2 monocytes and resident macrophages).

### Dendritic and myeloid-derived suppressor cells exhibit relatively small changes over disease progression

We identified 4 distinct populations of intrarenal dendritic cells that were infrequent before and after kidney injury in the 4 lupus mouse models (**Fig. 1B, C, E**). The most abundant population was cDC2 that displayed transcriptional similarities to previously described cDC2b cells (*44*) (**Fig. S10**). cDC1s were identified by *Clec9a*+, *Xcr1*+, *Tlr3*+ (**Fig. S1B**) and expressed functional gene sets for antigen processing and MHC class II presentation (**Fig. S10**). Ccr7+ DCs likely represented mreg-DCs (*45*) and expressed genes identified in tissue migratory cDCs (**Fig. S10**) that may facilitate transit to proximal lymph nodes (*46*). Plasmacytoid DCs expressed *Siglech*, *Tcf4*, *Il7r* (*47*) (**Fig. S10**) and were among the least frequent populations in all 4 strains (**Fig. 1C, E**). Finally, we identified a small cluster expressing genes that are associated with granulocytic myeloid-derived suppressor cells (*48*) (*49*) (**Fig. S1B**). All DC subsets exhibited relatively minor transcriptional changes from early to late disease consistent with the short renal half-life of these cells (*50*).

### Mice and humans have similar intrarenal myeloid subsets and gene programs in lupus kidneys

To determine whether patients have similar myeloid subsets we compared expressed myeloid transcripts from mouse myeloid to human cells from the kidneys of 155 unique patients with lupus nephritis and 30 healthy controls (profiled in a forthcoming larger study as part of the NIH AMP consortium). Kidney biopsies were processed for single cell RNA sequencing resulting in 23,819 high-quality intrarenal myeloid cells. To enable comparative analysis and identify human counterparts of the mouse renal myeloid subsets, we used Harmony (*51*) to integrate and embed the mouse and human single cell datasets into a common low dimensional space which removed effects of species, sample, and 10x technology. We then assigned mouse cluster identities to human cells by k-nearest neighbor label transfer in the Harmony embedding, and ran uniform manifold approximation and projection (UMAP) to visualize the embedding (**Fig. S11A, Fig. 4A**). We calculated confidence scores for each human cell after label transfer and found highest scores for DC1 and pDCs and lower scores for MDSCs (**Fig. S11B, C**), possibly due to cellular heterogeneity in humans not present in mice. Our analysis revealed that intrarenal mouse myeloid clusters have human counterparts that were present in similar proportions as in mice. Specifically, resident macrophages and NC1 monocytes were the most abundant populations followed by classical monocytes. Dendritic cells were less frequent, with cDC2 the most abundant of the DCs. NC2 monocytes were rare in patients. To better understand the transcriptional overlap between mouse and human we compared the log-fold change of differentially expressed genes from equivalent clusters relative to all other myeloid cells (**Fig. 4B-E**) and performed gene set enrichment analysis. Resident macrophages shared gene programs for phagocytosis and antigen processing and presentation (**Fig. 4B, right**). Mouse and human C1 monocytes expressed gene sets associated with transendothelial migration, cell adhesion, glycolysis and HIF-1 signaling, (**Fig. 4C, right**). C2 monocytes shared gene programs for oxidative phosphorylation, phagocytosis, and cholesterol homeostasis (**Fig. 4D, right**). NC1 monocytes from both species expressed genes that regulate the actin cytoskeleton, transendothelial migration, FcGR mediated phagocytosis (**Fig. 4E, right**). Our integrated analysis thus reveals equivalent cell types and gene programs across species.

**Figure 4.**
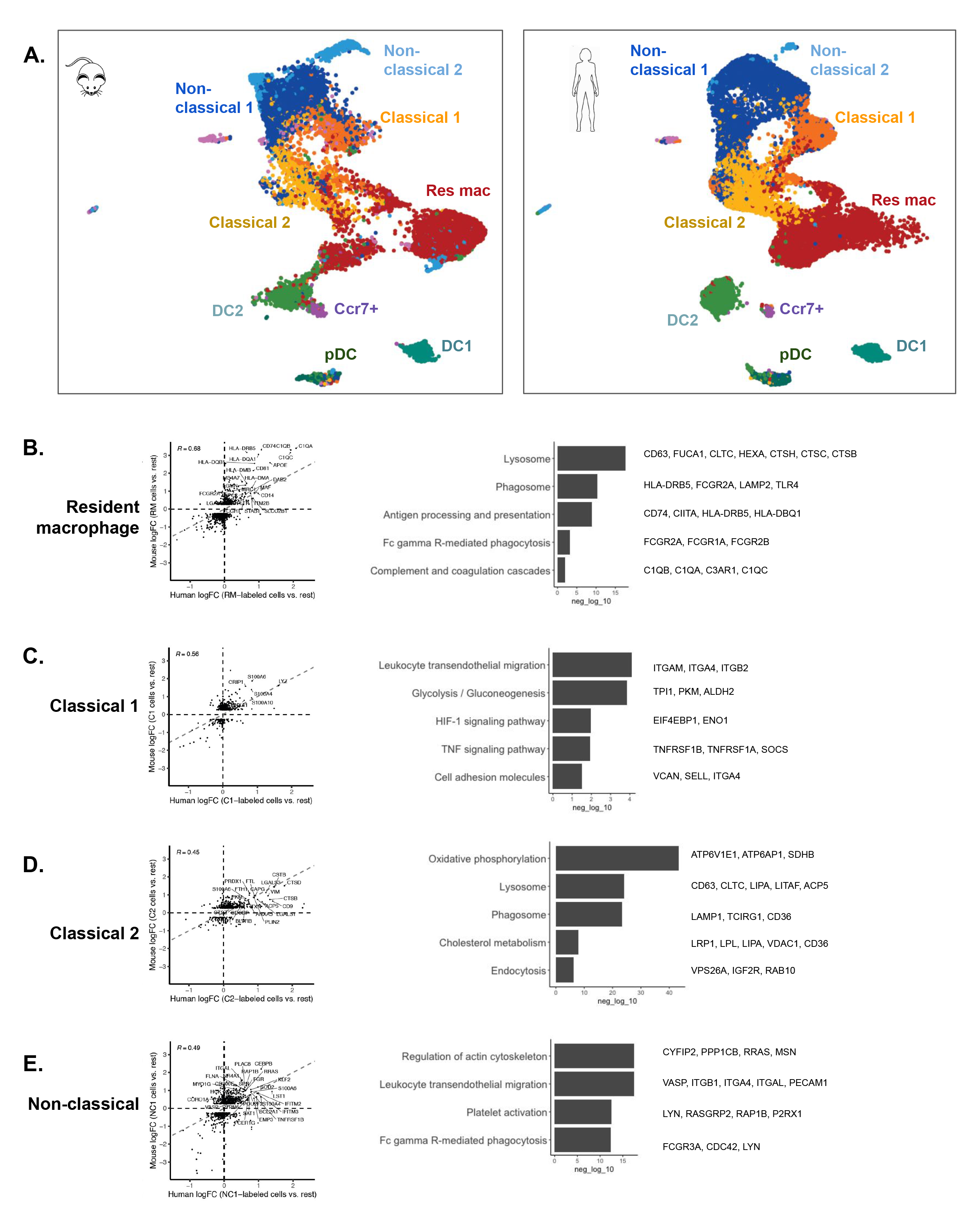
Integrated single cell analysis of kidney myeloid cells from 4 lupus mouse models and human patients (n= 155 with lupus nephritis, 30 healthy controls). **(A**) Split UMAP embeddings of integrated single cell analysis of myeloid cells colored by cell identities. Left: mouse. Right: human (identities mapped from mouse to human data by k-NN label transfer). (**B-E)** Left: scatter plots of cluster-specific marker genes from mice (y-axis) vs. humans (x-axis) comparing their expression (log-fold change) relative to other renal myeloid cells. Right: conserved gene sets enriched in KEGG analysis

### Human macrophages split into RM1 and RM0 subclusters comparable to those in mouse and their frequencies are associated with patient disease indices

Because the frequency of mouse RM1 decreased while RM0 increased with disease in mice (**Fig. 2A**), we examined changes in the equivalent cells in patients in relation to disease progression. We used Harmony to transfer mouse cluster identities to human cells as above. RM1 co-clustered with two myeloid populations (*LYVE1*+ and *LYVE1*-) present in healthy human controls (**Fig. 5A, B**). In contrast, RM0 co-clustered with those two plus an additional population that was only present in patients with lupus nephritis (**Fig. 5A, B**). Of note, *LYVE1*+ resident macrophages were not well represented in the 4 lupus mouse models. To examine the transcriptional relationship between human RM1 and human RM0, we applied PAGA to subclusters of human resident macrophages and identified the strongest relation between RM1 and RM0, consistent with a transition between these clusters (**Fig. S12**) as in mice (**Fig. 3C**). We compared differentially expressed genes between the equivalent mouse and human subsets to identify shared transcriptional changes during the putative RM1 transition to RM0. RM1 expressed cytokines and chemokines (*IL1B, CCL3, CCL31L1, CCL4, CCL4L2*) while RM0 expressed injury-associated genes *CD9, SPP1,* and *TREM2* as well as genes involved in lipid and cholesterol metabolism and lysosomal digestion (**Fig. 5C,D**). To determine the clinical significance of these macrophage subsets, we examined the correlations of RM1 or RM0 abundance (relative to all other RM subclusters) with two clinically important, multimodal disease indices (‘activity’ and ‘chronicity’) based on kidney biopsy histopathology (*52*). We found that RM1 frequency negatively correlated with the activity index (**Fig. 5E**) while RM0 frequency was positively correlated with chronicity that is associated with progression to end-stage renal disease (*53*) (**Fig. 5F**).

**Figure 5.**
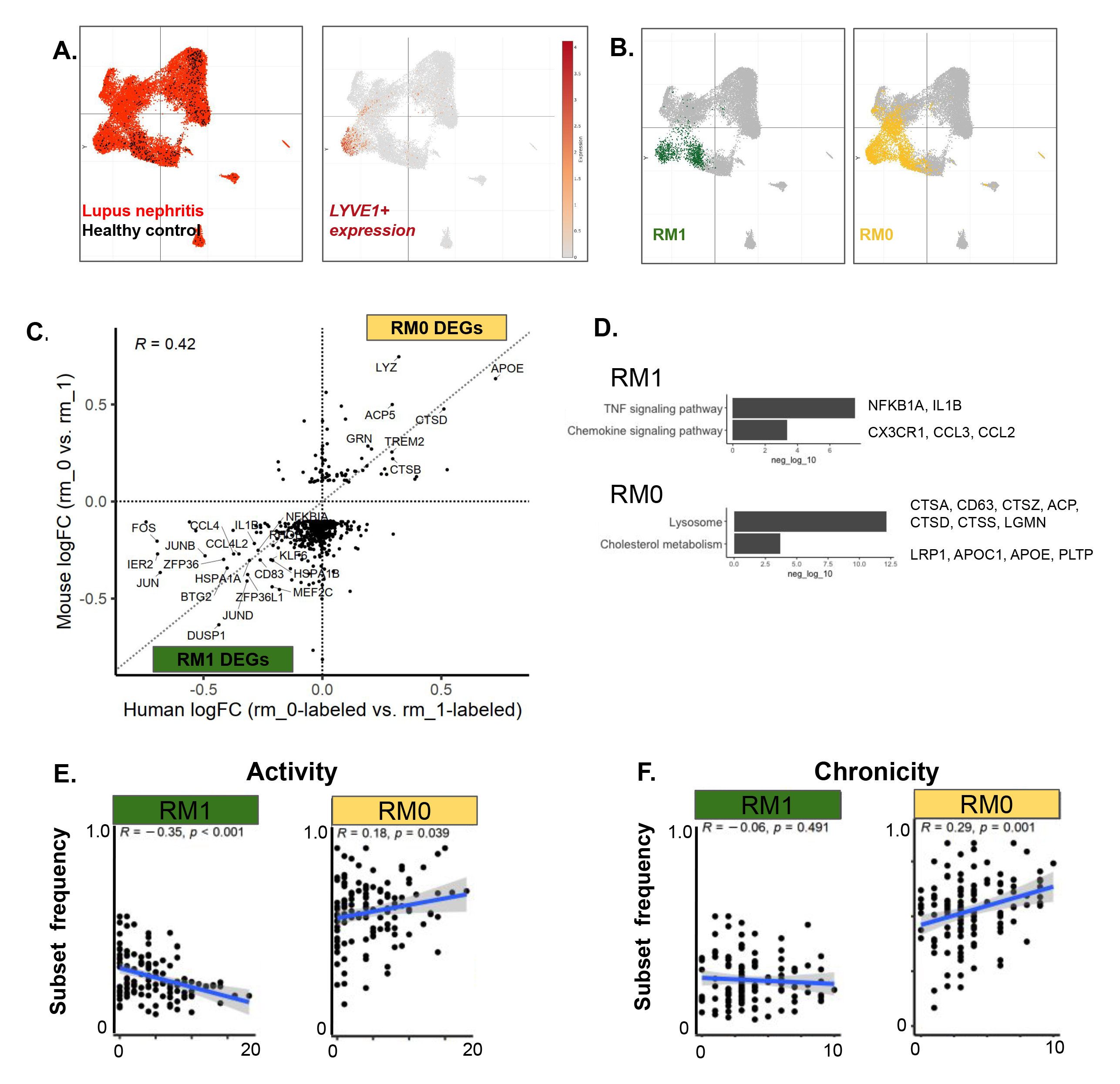
The frequency shift in RM1 and RM0 in humans is associated with histologic disease indices. **(A**) UMAP of renal myeloid cells from human patients colored by disease status (left): control in black; lupus nephritis in red; or by *LYVE1* expression (right). (**B**) UMAPs of RM1 and RM0 mouse identities mapped in human myeloid single cell data using k-nearest neighbor label transfer. (**C**) Scatter plot comparing the relative expression of RM1 and RM0 marker genes from mice (y-axis) vs. humans (x-axis). Left lower quadrant identifies conserved genes in RM1 and the right upper quadrant identifies conserved genes in RM0. (**D**) KEGG gene sets enriched in RM1 vs. RM0 from mice. (**E**) The frequency of human RM1 (left) or RM0 (right) per patient (n=123) relative to other RM subsets (y-axis) vs. the NIH Activity Index (x-axis). RM1 is negatively associated with the Activity Index and RM0 is positively associated. (**F**) The frequency of human RM1 (left) or RM0 (right) per patient (n=123) relative to other RM subsets (y-axis) vs. the NIH Chronicity Index (x-axis). RM1 has no association and RM0 is positively associated with the Chronicity Index.

### Human classical monocytes are associated with patient disease indices

Because classical 2 monocytes expanded in nephritic mouse kidneys (**Fig. 1D, E, Fig. 3A, C, Fig. S3A, B**) we further characterized the equivalent human population. C2 monocytes were infrequent in healthy kidney donors (**Fig. 6A, Fig. S13**). Relative to C1, C2 cells expressed *CD9, SPP1, APOE, FABP5 GPNMB,* and *TREM2* (**Fig. 6B**) that were previously reported in monocytes associated with tissue inflammation and fibrosis in multiple organs in humans and mice (*22*). We note heterogeneous expression of these markers across the human C2 cluster (**Fig. S14**), suggesting that C2 can be further subclustered. To test whether C2 might derive from C1 as we observed in mice (**Fig. 2F**), we again applied PAGA to human myeloid clusters labeled by their mouse identities. This revealed high transcriptional similarity between C1 and C2 (**Fig. S15)**. C2 may transition from infiltrating C1 in kidneys given our prior results that C2 was infrequent in the blood of patients with lupus nephritis (*7*). In patients with lupus nephritis, C2 cells were present and highly correlated with histopathologic activity but not with chronicity (**Figs. 6C, D**).

**Figure 6.**
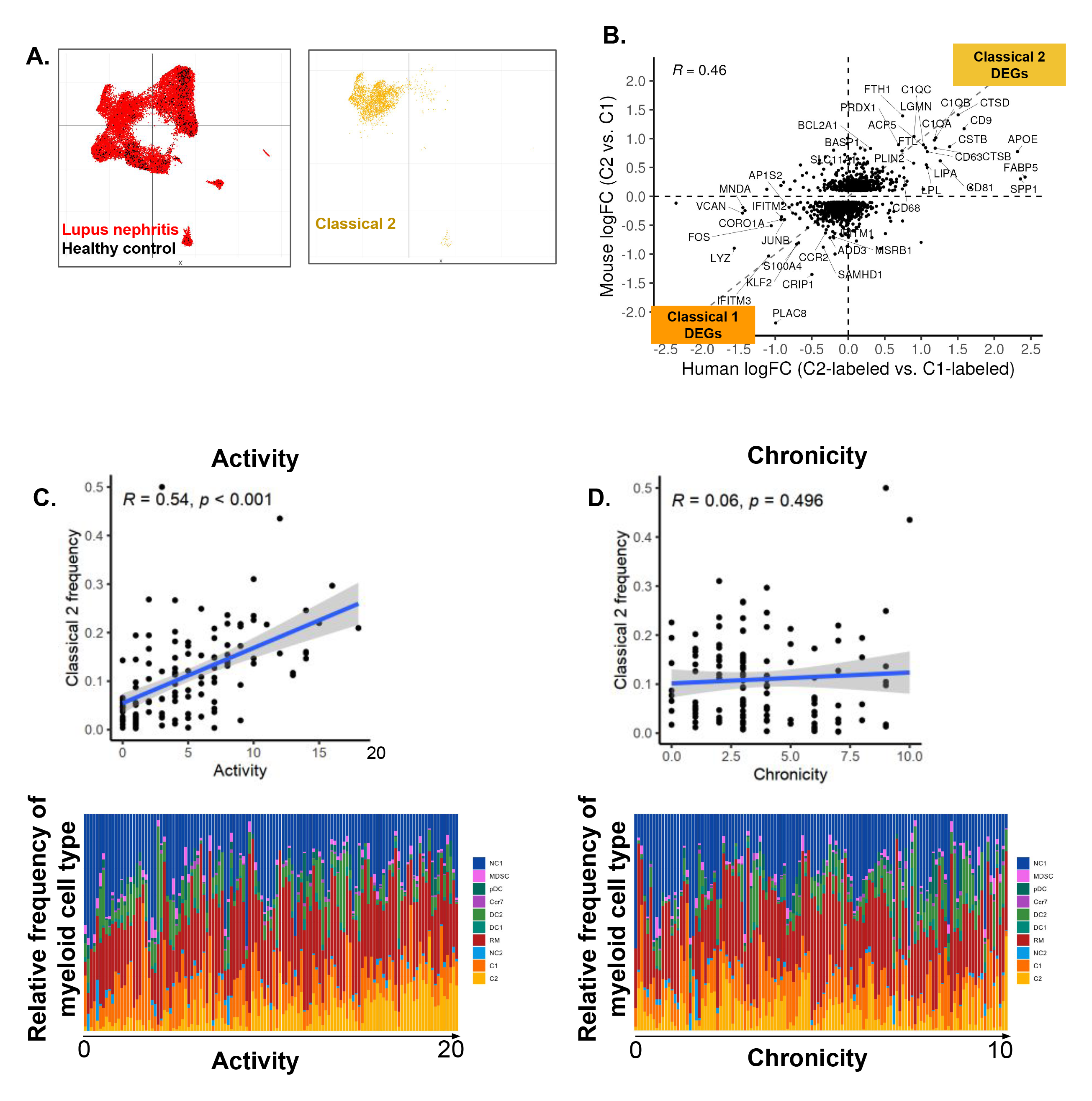
Classical 2 monocytes in humans are associated with histopathologic kidney injury in lupus nephritis. **(A**) Left: UMAP of renal myeloid cells from human patients colored by disease status: healthy (black); lupus nephritis (red). Right: Classical 2 monocyte mouse identities mapped onto the UMAP embedding of human myeloid single cell data using k-nearest neighbor label transfer. (**B)** Scatter plots comparing the expression of marker genes from Classical 1 and Classical 2 in mice (y-axis) vs. humans (x-axis). Left lower quadrant identifies conserved genes in Classical 1, and right upper quadrant identifies conserved genes in Classical 2. (**C**) The frequency of human Classical 2 monocytes per patient (n=123) relative to other myeloid subsets (y-axis) vs. the NIH Activity Index (x-axis). Classical 2 monocytes are positively associated with the Activity Index. Top: Scatter plot. Bottom: Normalized bar plots comparing frequencies of myeloid subsets from each patient by scRNA-seq droplet proportions. (**D**) The frequency of human Classical 2 monocytes per patient (n=123) relative to other myeloid subsets (y-axis) vs. the NIH Chronicity Index (x-axis). Classical 2 monocytes are not associated with the Chronicity Index. Top: Scatter plot. Bottom: Normalized bar plots comparing frequencies of myeloid subsets from each patient by scRNA-seq droplet proportions.

### Human non-classical monocytes are inversely associated with patient disease indices

Non-classical monocytes also play a role in mouse lupus nephritis by triggering an immune response that damages the glomerular endothelium upon exposure to TLR7 agonists (*10*) (*24*). Surprisingly, the frequency of NC1 monocytes in patients was inversely correlated with both activity and chronicity histologic indices in patients (**Fig. 6B bottom and 6C bottom, Fig. S16A, B**).

### Human classical 2 monocytes are enriched in a subset of patients with class III and IV lupus nephritis

Because C2 monocytes positively correlated with the activity index that primarily reflects glomerular histology (*52*), we tested whether C2 was associated with different forms of glomerular injury (class III/IV biopsies are characterized by both glomerular cellular hyperproliferation and immune complex deposition at subendothelial and class V by immune complex deposition at subepithelial sites). Relative to other myeloid subsets, C2 monocytes were enriched in class IV and to a lesser extent in class III kidney biopsies, with or without concomitant class V lesions, when compared to biopsies with pure class V lesions or control (**Fig. 7A**) or to patients with acute and chronic kidney disease (**Fig. 7B, S17A, B**). In contrast, RM0 (and all other myeloid subclusters) was present at similar frequencies across all histologic classes (**Fig. S18B**) as well as acute and chronic kidney disease (**Fig. S18D, S19**). We conclude that C2 monocytes are relatively specific to class III and IV lupus nephritis, whereas RM1 and RM0 are shared across histologic classes III, IV, V and acute and chronic kidney disease.

**Figure 7.**
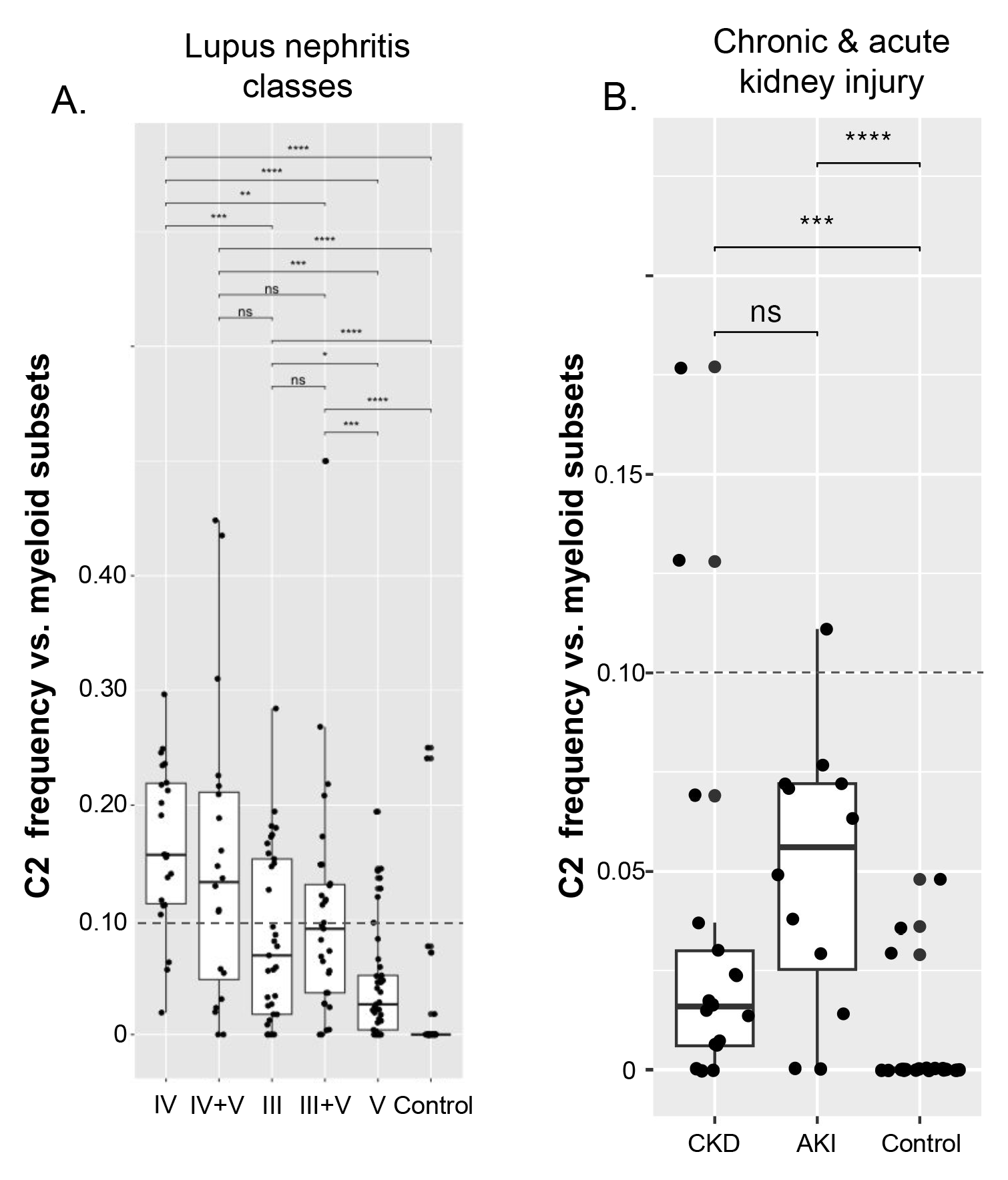
Classical 2 monocytes are enriched in patient biopsies with class III and IV histologic lesions compared to class V alone. Boxplots depicting the distribution of patients with class IV, IV+V, III, III+V, or healthy controls and the frequency of Classical 2 relative to other myeloid subsets (**A**) or chronic kidney disease and acute kidney injury (**B).** Kruskal-Wallis test comparing means (ns: p > 0.05; *: p < 0.05; **: p < 0.01; ***: p < 0.001; ****: p < 0.0001).

### *In situ* localization of comparable myeloid cell states in human lupus nephritis kidney sections

To localize myeloid subsets in kidney sections from patients with lupus nephritis, we again employed RNA FISH probes to label cells from the myeloid lineage (*CD68, CSF1R*) and monocyte and macrophage subsets using 2-3 cluster-specific genes from our scRNA-seq analysis (**Fig. S20**). Classical 1 monocytes were rare in clinical nephritis. When present, they were in the glomerulus rather than the tubulointerstitium or the immune infiltrate (**Fig. S21A, Fig. S22A**). Classical 2 monocytes concentrated in glomerular and peri-glomerular sites (**Fig. S21B**) and also in immune infiltrates (**Fig. S22B**). Non-classical 1 monocytes localized to glomerular sites but were rare in the tubulointerstitium (**Fig. S21C, Fig. S22C**). Finally, resident macrophages localized to the tubulointerstitium where they aggregated around nephritic glomeruli and formed an interspersed network throughout the interstitium (**Fig. S21D, Fig. S22D, left**) but were also close to interstitial immune cell aggregates in the diseased tissue (**Fig. S22, right**), consistent with previous work in mice (*41*) (*42*) (*43*). Our data reveal that transcriptionally comparable myeloid subsets occupy similar regions in kidneys from humans and mice with lupus nephritis where they express similar functional gene programs.

## DISCUSSION

This is the most extensive study to date of intrarenal myeloid states in lupus nephritis. It encompasses cellular definitions that are mapped between mice and humans, localization to renal microenvironments, and inferences about the origin of cellular states. This atlas is based on 4 mouse models of lupus nephritis with different genetic backgrounds and pathogenic drivers at preclinical and clinical stages of disease and 155 adult patients with diverse ethnic/racial backgrounds with different stages and histologic classes of disease. Within our cross-species comparison, we identified classical 2 (C2) monocytes and resident macrophage (RM) states that expressed a conserved set of injury-associated genes as well as programs for debris clearance, lipid metabolism, and tissue repair that correlate with advanced disease, pointing to conserved functional pathways associated with clinically relevant kidney damage in humans. This atlas can be used as a resource and a starting point for mechanistic and *in vivo* studies.

C2 monocytes in humans were strongly associated with proliferative lupus nephritis (the most severe form of disease) and expanded with disease progression in kidneys from all 4 lupus mouse models. These cells occupied both glomerular and interstitial niches including renal immune aggregates that are present in hypoxic areas (*54*). We hypothesize that C2 monocytes differentiate from infiltrating C1 monocytes based on our PAGA analysis and their low representation in the blood in mice. Differentiation of C1 to C2 may be induced by distinct types of kidney injury that are characteristic of class III/IV glomeruli, like hyper-proliferative glomerular cells, Fc receptor mediated activation by glomerular immune complexes (deposited between the vascular endothelium and the basement membrane) (*52*) (*55*), and exposure to DAMPs from injured cells or circulating inflammatory mediators. These factors could enable tissue infiltration (*56*) and/or provide differentiation cues. A separate or overlapping set of factors in the interstitium may also induce differentiation of C1 to C2, although their identities are less clear since the histologic differences in the interstitium between classes III/IV vs. V are less pronounced than the glomerulus. Of note, cells similar to C2 monocytes have been derived *in vitro* from human CD14+ monocytes in response to combination GM-CSF, IL-17A, and TGF-B1 (*22*), or to pathogenic lipids (*57*), and after 72 hours of culture (possibly from apoptotic cells in the cultures) (*58*). Thus, the C2 phenotype may represent a conserved injury response that can be induced by unique factors from the glomerulus or the interstitium.

C2 monocytes in both species expressed functional gene sets for phagocytosis, lysosomal functions, and cholesterol metabolism as well as an injury-associated gene set (*CD9, SPP1, APOE, FABP5, GPNMB, PLA2G7* and *TREM2).* Monocytes with similar functional pathways and genes were previously reported in association with fibrosis and tissue inflammation in models of injury in multiple organs (but not in a predictive model of macrophage activation derived using acute infectious stimuli and applied to 12 mouse tissues (*59*)). C2-like macrophages localized to inflamed and scarred regions in cirrhotic livers (scar associated macrophages - SAMs) (*60*), were found in injured lung (*61*), atherosclerotic arteries (*62*) and inflamed brain (damage associated macrophages - DAMs) (*21*), and arose in adipose tissue from mice fed a high fat diet (lipid associated macrophages - LAMs) (*23*). Our data suggest that we have identified the renal counterpart of these cells in humans and mice. C2-like macrophages have been proposed to promote fibrosis in liver (*22*) (*60*) and lung (*22*). However it has been difficult to associate pathogenic and profibrotic functions of these cells with the expression of the injury-associated genes. *Spp1*-deficiency promoted liver fibrosis in NASH (*63*) and *Fabp5-*deficiency induced M2 polarization (*64*) through alterations in fatty acid metabolism. *Gpnmb* reduced macrophage inflammatory functions in obese mice (*65*). *Trem2*-deficient SAMs were pro-fibrotic and pro-inflammatory (*66*) and *Trem2*-deficient mice failed to generate LAMs resulting in defective adipocyte clearance and obesity (*23*). The role of *Pla2g7* is less clear since it has been associated with both pro-and anti-inflammatory macrophage functions (*67*) (*58*). Collectively, these injury-associated genes appear to modulate fibrosis and inflammation in tissue-and context-specific manners. In the human kidney C2 cluster, expression of the injury-associated genes is heterogeneous, suggesting the presence of subclusters that could differentially modulate repair and fibrosis (*22*).

Our finding that C2 monocytes were more abundant in biopsies with class III and IV lesions compared to class V and other forms of kidney disease may have clinical implications. First, because C2 are the major myeloid subset in the urine of LN patients (*7*), they may be useful to distinguish between patients with class III/IV lupus nephritis from class V and other forms of acute and chronic kidney disease. Second it could be possible to monitor the response of LN patients to treatment by measuring urine C2. Proteomic analyses of urine from patients in the AMP cohort revealed that the decline of urinary macrophage markers CD163 and CD206 outperformed the clinical standard (proteinuria) at predicting 1-year response to therapy (https://ard.bmj.com/content/82/Suppl_1/138.1). Further investigations will be required to establish the relationship and the specificity of urine C2 cells and associated proteins to disease activity and therapeutic responses.

Mouse and human kidney resident macrophages (RM) are normally positioned in the tubular interstitium between the tubules and peritubular capillaries where they monitor trans-endothelial transport and may initiate an inflammatory response shortly after ingesting circulating immune complexes (*11*), particularly those containing endosomal TLR ligands. In contrast to C2, RM subset frequency did not differ across histologic classes of lupus nephritis. RM was dominated by RM1 and RM0 sub clusters in both species. In mice, RM1 frequency shifted toward RM0 with disease progression. We observed a similar shift in humans that was positively associated with the NIH chronicity index that reflects inflammatory and fibrotic changes in the tubulointerstitium and glomeruli and portends poor clinical outcomes (*53*). We acknowledge that RM1/RM0 frequencies were not significantly different in lupus nephritis patients vs. controls. This may be due to low numbers of RMs in most healthy controls in our cohort or to possible modulation of the RM0 phenotype with treatment (*50*). We also note that RM0 co-clustered with RM1 plus an additional population present only in patients with lupus nephritis, supporting a shift from RM1 to RM0 associated with chronicity in humans. We hypothesize that RM1 and RM0 are related by differentiation based on our PAGA analyses. RM0 could originate in response to ingestion of immune complexes, tissue debris or exposure to DAMPs released as a result of chronic local or systemic inflammation. RMs could also derive from local cellular proliferation of the small RM population in mice that expressed *Mki67* (*68, 69*) but we did not observe this in humans. Blood precursors could also enter inflamed kidneys and differentiate to resident macrophages during kidney injury as reported in parabiosis studies (*70*). RM0 expressed gene programs for lysosomal functions, cholesterol metabolism and several genes from the C2 injury-associated gene set. Thus, similar functional gene programs and injury-associated genes were present in both resident macrophages and infiltrating monocytes in lupus kidneys in both species. This is not without precedent, as resident microglia in Alzheimer’s disease in mice and humans (*21*) and infiltrating monocytes in human liver fibrosis (*60*) that express similar genes have been found.

Non-classical (NC) 1 monocytes are known to patrol endothelial lumens in glomeruli (*10*). In our human cohort, NCs were present in kidney biopsies from healthy controls and those with low activity or low chronicity. Surprisingly, NC1 frequency decreased with increasing activity and chronicity indices, possibly due to differentiation into classical 2 monocytes based on our PAGA analyses. In contrast, mouse intrarenal NC1s increased with disease progression that likely reflect their peripheral expansion, a feature of active disease that is not observed in humans (*71*). NC1 from both species expressed genes that may contribute to lupus nephritis pathogenesis. *ITGAL* promotes homeostatic patrolling behavior on endothelium, *CX3CR1* interacts with endothelial *CX3CL1* in the presence of a nucleic acid danger signal, and *CCL3* that may recruit tissue-damaging neutrophils in both species. In knockout studies, these genes were critical for non-classical monocytes to accumulate in the glomeruli where exposure to a TLR7 agonist increases their endothelial retention and triggers an inflammatory cascade that damages and facilitates removal of endothelial cells (*10*) (*24*) (*72*). Thus, conserved gene expression suggests that mechanisms first identified in mouse NC likely impact human lupus nephritis. We also identified in mice (but not humans) a distinct NC2 monocyte that localized to glomeruli characterized by expression of *Itgal* and *Nr4a1* like NC1 but with additional pro-and anti-inflammatory gene sets. The role of NC2 cells is still unknown.

There are several notable limitations to our study. First, we were not able to identify perfectly discriminating markers to stain myeloid cell subsets in kidney sections, and further discovery and application of markers will be needed. Second, mapping mouse cell identities to human cells could be imperfect (as we observed in the confidence scores we calculated for each cell). Third, our analysis of cell trajectories infers differentiation pathways between cell types but only lineage studies can prove clonal relationships. Fourth, while we identify conserved disease-associated myeloid subsets, we have not yet developed the tools needed to study the impact of individual subsets in lupus nephritis disease in mice. Fifth, our study included single biopsies from each patient regardless of treatment. Therefore, we could not compare timepoints in the same patient nor isolate the impact of different treatments across patients. Lastly, human biopsies were primarily collected from the kidney cortex while mouse data were generated from the whole kidney. Nevertheless, our analyses identified comparable intrarenal myeloid subsets with conserved alterations in classical monocytes and resident macrophages associated with advanced disease across species.

By comparing intrarenal myeloid populations in mice and humans with lupus nephritis we identified congruent cell types and pathways, including convergent programs for debris clearance, lipid metabolism, and tissue repair in resident macrophages and classical 2 monocytes that are associated with advanced disease in both species. These findings provide a basis for testing mechanistic hypotheses derived from human tissues in mouse models, and a map to develop tools that manipulate these subsets and their genes to study the role of human risk variants in their function, determine their lineage relationships and discover their impact on disease and response to therapies.

## METHODS

### Mouse colonies, myeloid cell isolation, and single cell genomic library generation

**Mice.** Males from Tlr7-overexpression (B6.Sle1.Yaa and NZW/BXSB) and females from lymphoid-expansion (NZB/W and MRL/lpr) mice were followed clinically as previously described (*42*) (*73*). Briefly, urine was tested every other week for proteinuria by dipstick (Multistick; Fisher Scientific) or BUN by bleeding. Once fixed proteinuria of 300 mg/dl on two occasions 24-h apart appeared plus elevated proteinuria of BUN > 30 mg/dL for 2 weeks, mice were considered to have clinical nephritis. We analyzed mice with preclinical or clinical nephritis before the onset of terminal renal failure. All experiments were approved by the IACUC of the Feinstein Institute for Medical Research.

### Murine kidney dissociation and single cell suspension

Under terminal anesthesia, mice were perfused by gentle intracardiac injection of 10 ml prewarmed (37°C) PBS 1x and kidneys were harvested. To obtain single cell suspensions, kidneys were then incubated for 15 minutes in DMEM containing 1 mg/ml Collagenase D (Roche), 100U/ml DNAse I (Sigma), 0.25mg/ml Liberase (Invitrogen) and 3% heat-inactivated fetal bovine serum (FBS, Invitrogen) at 37°C. Tissues were mechanically dissociated and then passed through a 100 μm cell strainer (BD). This single cell suspension was finally enriched for CD45+ cells using Mouse CD45 MicroBeads (Miltenyi) and LS magnetic Columns (Miltenyi) for FACS sorting.

### Murine blood processing

Prior to termination, whole blood from mice was collected in heparinized tubes that was then added to Pharma Lysis buffer (BD) for 20 minutes at room temperature. PBMCs in lysed blood were then washed twice with sterile PBS and placed in FACS buffer for downstream applications. For flow cytometric analysis, PBMCs were blocked with rat serum and incubated with fluorescent antibodies in a staining buffer (BD). For 10x, PBMCs were sorted based on size to exclude doublets, dead cells, cellular debris, or any RBC remnants. Sorting was performed on BD FACS Aria II.

### Murine single cell RNA-seq library preparation and sequencing

Single cell suspensions of myeloid or immune cells from the kidneys or blood from 2 to 4 age-and strain-matched mice with preclinical or clinical nephritis were pooled and washed in sterile PBS. Up to 10,000 live cells were loaded into separate 10x channels for single cell RNAseq on the Chromium platform (10x Genomics). DNA amplification and library construction were carried out according to the manufacturer’s instructions. The purified libraries were quantified and sequenced according to manufacturer’s guidelines using Nexteq or Novaseq S1 platforms (Illumina).

### Human kidney biopsy collection, dissociation, and single cell genomic library generation

#### Biopsy collection

As part of the Accelerating Medicines Partnership (AMP) RA/SLE Phase 2 consortium patients >16 years old undergoing a clinically indicated kidney biopsy to evaluate proteinuria (urine protein to creatinine ratio >0.5) were enrolled if they met sufficient criteria of systemic lupus erythematosus diagnosis based on the revised American College of Rheumatology or the Systemic Lupus Erythematosus International Collaborating Clinics classification criteria (*74*), (*75*). Patients were excluded if they had a history of kidney transplant, rituximab within 6 months of biopsy, or were pregnant. All patients provided written informed consent.

#### Histologic scoring of human lupus nephritis kidney sections

Scoring was performed centrally by two board-certified pathologists (JH and DD). The NIH “Activity” index represents ongoing kidney damage that is scored by the extent of endocapillary hypercellularity, karyorrhexis/neutrophils, fibrinoid necrosis, subendothelial deposits, luminal thrombus, cellular and fibrocellular crescent formation, and interstitial inflammation (*52*). “Chronicity” represents kidney damage that is often irreversible and is scored by the extent of sclerosis in the glomeruli, fibrotic crescents, tubular atrophy, and interstitial fibrosis.

#### Human kidney biopsy dissociation and single cell suspension

Human kidney biopsies were cryopreserved, thawed and dissociated as described (*7*), with modifications. Briefly, samples were dissociated enzymatically with 0.5mg/mL Liberase TL (Roche) in DMEM/F12 (Corning) for 12 minutes at 37C, and then mechanically with a cell strainer pestle (CELLTREAT) on a 70 um strainer (Miltenyi). Cells were washed with cold RPMI (ThermoFisher) supplemented with 10 mM HEPES and 0.04% BSA (Millipore Sigma) and resuspended in 50uL of the same medium. The samples were then filtered through a 40 um strainer (Pluriselect), and after counting by Trypan exclusion, up to 10,000 cells were loaded onto Chromium microfluidic chip for single cell transcriptomics profiling using the 3’ V3 kit (10x Genomics). Sequencing libraries were produced according to the manufacturer’s instructions. Prior to sequencing, DNA fragments derived from mitochondrial transcripts were digested using Depletion of Abundant Sequences by Hybridization (DASH) (*76*), with modifications to optimize for 3’ 10x libraries. Following DASH, libraries were sequenced on a Novaseq S4 (Illumina) according to the manufacturer’s guidelines.

### Bioinformatic Analysis

#### Processing of mouse scRNA-seq data

Single cell suspensions of intrarenal or peripheral myeloid or immune cells from 2 to 4 age-and strain-matched mice with preclinical or clinical nephritis were prepared using 10x 3’ or 5’ chemistry. In total, the mouse data set included 20,299 myeloid cells after quality control (described below). Fastq files were aligned to the mm10 genome reference with Cellranger v6.0.1 All downstream analysis was performed using R v4.1 and Seurat v.4. The B6.Sle1.Yaa and NZBW strains were each multiplexed with four samples and required demultiplexing with Seurat’s HTODemux function. In each case, cells assigned to the one blood sample were removed, leaving only cells from three kidney samples. Cells classified as inter-sample doublets were also removed. We next performed the following analysis steps separately on all four mouse strains: (i) Ambient RNA was estimated and removed using the R package Soupx v.1.6. (ii) Cells with <500 or >5,000 genes or >5% mitochondrial RNA expression were removed. (iii) A standard Seurat pipeline was used with default parameters for these steps: NormalizeData, FindVariableFeatures, ScaleData, RunPCA, FindNeighbors, FindClusters and FindAllMarkers. (iv) Clusters were manually inspected for various markers and quality metrics, and several clusters representing low quality, non-myeloid and proliferating cells and doublets were removed. Seurat objects from each of the four strains were then merged into one object, to which the standard Seurat pipeline was applied with the following modifications:

A. Rather than calculating variable features across the four strains, which would overemphasize differences between strains, we used this method:

Step 1: Calculate variable features on each strain subset separately, and rather than returning the top 2000, return all.
Step 2: Rank each of the four lists from most variable (rank = 1) to least variable (rank = # of features).
Step 3: For each feature, calculate the average of the 4 strain ranks.
Step 4: Take the top 2001 features with lowest average rank.
B. After performing PCA, we performed batch correction with the R package Harmony v0.1.1 (*51*), harmonizing on strain with theta=2. Subsequent steps (FindNeighbors and RunUMAP) used Harmony components instead of principal components.

Finally, differentially expressed genes were identified using Seurat’s FindAllMarkers and the resulting clusters were manually annotated based on differentially expressed canonical markers as described in the main text.

#### Processing of human scRNA-seq data

11 samples were prepared using 10x v2 chemistry, and the rest used 10x v3 chemistry. In total, the human dataset included 23,819 myeloid cells. Single-cell RNA-sequencing data was aligned using the 10x Genomics Cell Ranger pipeline with the GRCh38 reference transcriptome. Cells with at least 500 detected genes, 1000 UMIs and less than 3% reads mapped to mitochondrial genes (following DASH) were kept for subsequent analysis. Scrublet (*77*) was used to remove suspected doublets, such that the expected doublet rate was calculated based on the number of cells loaded, and parameter values were set to min_counts=2, min_cells=3, min_gene_variability_pctl=90, and n_prin_comps=20.

#### Translating mouse gene symbols to human gene symbols

To integrate the mouse and human expression data, we converted the mouse gene symbols to human gene symbols by mapping between mouse and human gene orthologs. We obtained the ortholog mapping using the ‘biomaRt’ R package (v2.46.3), mapping mgi_symbol from the mmusculus_gene_ensembl database to hgnc_symbol from the hsapiens_gene_ensembl database, using the dec2021.archive version of Ensembl. We added additional ortholog pairs from HomoloGene (https://ftp.ncbi.nih.gov/pub/HomoloGene/build68/) to obtain a total of 19,055 ortholog pairs (representing D=17,798 unique human genes and d=18,230 unique mouse genes). We represented this mapping as a matrix with human genes as rows, mouse genes as columns, and values in {0,1} denoting whether a mouse gene maps to a human gene. The vast majority of mouse genes (97% = 17,701/18,230) had one-to-one mappings. To handle the one-to-many mappings, we normalized each column to sum to one to create a count-preserving probabilistic map from mouse to human genes M ∈ R^D×d^. We then applied the mapping matrix to the original mouse expression matrix (U_mouse_) to obtain the “humanized” mouse expression: U_humanized_= MU_mouse_. In this way, the UMI counts for a given mouse gene were uniformly distributed among the matching human symbols for one-to-many mappings. For any human orthologs that were missing in the mouse expression data, we filled in the expression with zeroes.

#### Mouse and human integration and label transfer

Starting from the human expression matrix and humanized mouse expression matrix, we performed log(CP10K+1)-normalization on each cell. We subset by the set of 16,957 genes overlapping between the two species and concatenated the cells from both species into a combined matrix. We calculated the top 1,500 variable genes within each species separately using the variance-stabilizing transform method in Seurat. After merging the two variable gene sets, we removed ribosomal, mitochondrial, and cell cycle (g2m.genes and s.genes in Seurat v4.1.0) genes, resulting in 2,345 variable genes used for subsequent dimensionality reduction. We scaled the data and performed PCA using the ‘Seurat’ R package (v4.1.0), calculating the top 20 PCs. We then removed the effect of species, sample, and technology (10x version) using Harmony (v0.1.0) with theta = 2.5, 0.5, and 0.5, respectively. This resulted in an integrated low-dimensional embedding of cells from both species. We ran uniform manifold approximation and projection (UMAP) using the uwot R package (v0.1.11) to visualize the embedding. To transfer the cell cluster labels defined in the mouse dataset to the human cells, we predicted the mouse label for each human cell by taking the majority vote of the 5 closest mouse cell neighbors in the Harmony embedding based on Euclidean distance (with ties broken at random), using the ‘class’ (v7.3-20) R package.

#### Comparing differentially expressed genes across species

To compare differentially expressed genes (DEGs) for each cluster across species, we used the ‘wilcoxauc’ function from the presto (v1.0.0) R package to calculate DEGs within each species separately (each cluster relative to cells from all other clusters). Human clusters were defined using the label transfer from mouse cells. We then compared the logFC of each DEG in each mouse cluster to the logFC of the same gene in each human cluster, limiting the comparison to significant DEGs in mice (Benjamini-Hochberg adjusted *P*-value < 0.01).

#### Inferred differentiation of myeloid clusters

To identify transcriptional relatedness between clusters and infer differentiation in our scRNA-seq data, we used Partition-based Graph Abstraction (PAGA) based on the *k*-NN graph above (*20*). To move from Seurat to the Anndata format supported in the Scanpy and Scvelo Python libraries, the SeuratDisk package in R was used to convert Seurat objects to the h5ad compressed format. Metadata labels are not carried over during this conversion process, so all labels were exported and loaded separately. A diffusion map and a diffusion pseudotime was generated for each dataset for the PAGA analysis. For each analysis, the classical 1 myeloid subset was used as the starting cluster. All connections between clusters in the PAGA plots were given a connectivity score between 0 and 1. UMAP overlay and circular graphs were generated using these connectivity scores.

### In situ localization of cellular states identified by scRNA-seq

#### RNAscope in situ hybridization with co-immunostaining

Lupus nephritis biopsies were identified using the Brigham and Women’s Hospital (BWH) Lupus Center Registry of validated SLE cases (*78*) fulfilling 1997 ACR criteria for SLE Classification (*74*). 3-micron thick sections were cut from formalin-fixed paraffin-embedded mouse or human kidney blocks onto Permaflex Plus slides and baked at 60C for one hour before use. Mixed RNAscope (Advanced Cell Diagnostics)/antibody antigen retrieval and staining with Opal (Akoya Biosciences) fluorophores was performed manually following the RNAscope Multiplex Fluorescent v2 Assay combined with the immunofluorescence protocol (322818-TN) and the RNAscope 4-plex Ancillary Kit for Multiplex Fluorescent Reagent Kit v2 Technical Note (323120-TN). All fluorophores were diluted at 1:750. Slides were coverslipped (Fisher Scientific) and mounted with ProLong Gold Antifade Mountant with DAPI (ThermoFisher). Stained slides were imaged using a Vectra Polaris microscope simultaneously with positive and negative control slides using identical image acquisition settings.

#### Image analysis and ROI selection

Multi-color raw images were acquired with the Vectra Polaris. TIFFs from single fields of view were then stitched together fused into a single multi-layer pyramidal TIFF in Halo software (Indica Labs) that were unmixed using inForm software (Akoya Biosciences) utilizing an algorithm based on a library of fluorescence spectra generated from control slides labeled with single fluorophores. Signal intensity thresholding was set according to levels of background in situ hybridization fluorescence in negative control slides. Morphologic features in both mice and humans were identified based on morphological features from pathologists (JH) and included the glomeruli, the tubulointerstitium, lymphoid aggregates, and blood vessels. In mice, glomeruli were also identified using antibody co-stain (Nphs2) targeting a podocyte-specific epitope.

## SUPPLEMENTAL FIGURES

**Sup. Fig. 1.**
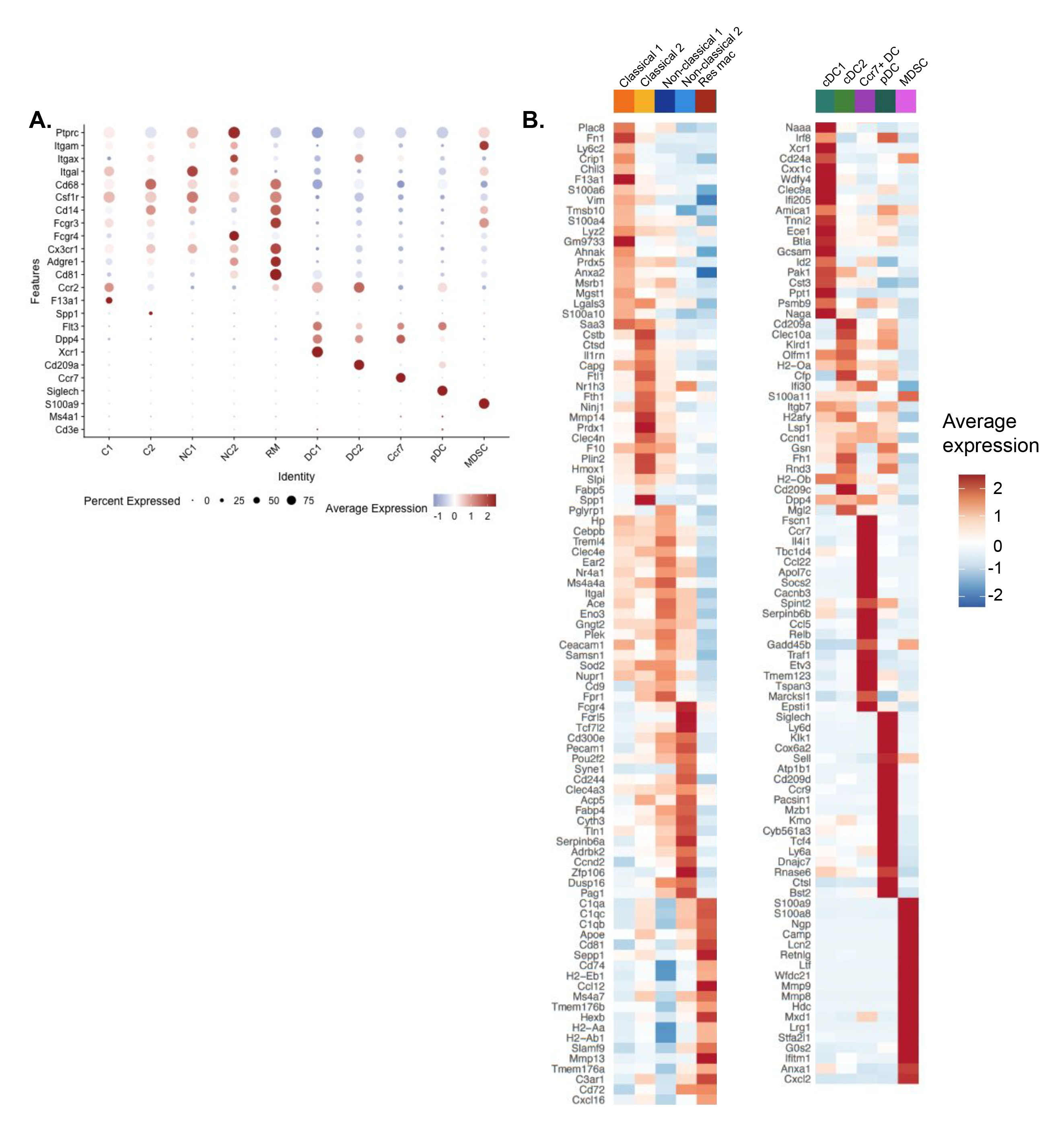
Identification of intrarenal mouse myeloid subsets by lineage-specific genes. **A**. Dot plot depicts markers used to discriminate monocytes, macrophages, dendritic and T and B-cells. Color scheme is based on z-score, from −1 (blue) to 2.3 (red). **B**. Heatmap shows distribution of scaled expression of top 20 discriminative genes for each myeloid cluster in Fig. 1C. Color scheme is based on z-score, from −2.2 (blue) to 2.2 (red).

**Sup. Fig. 2.**
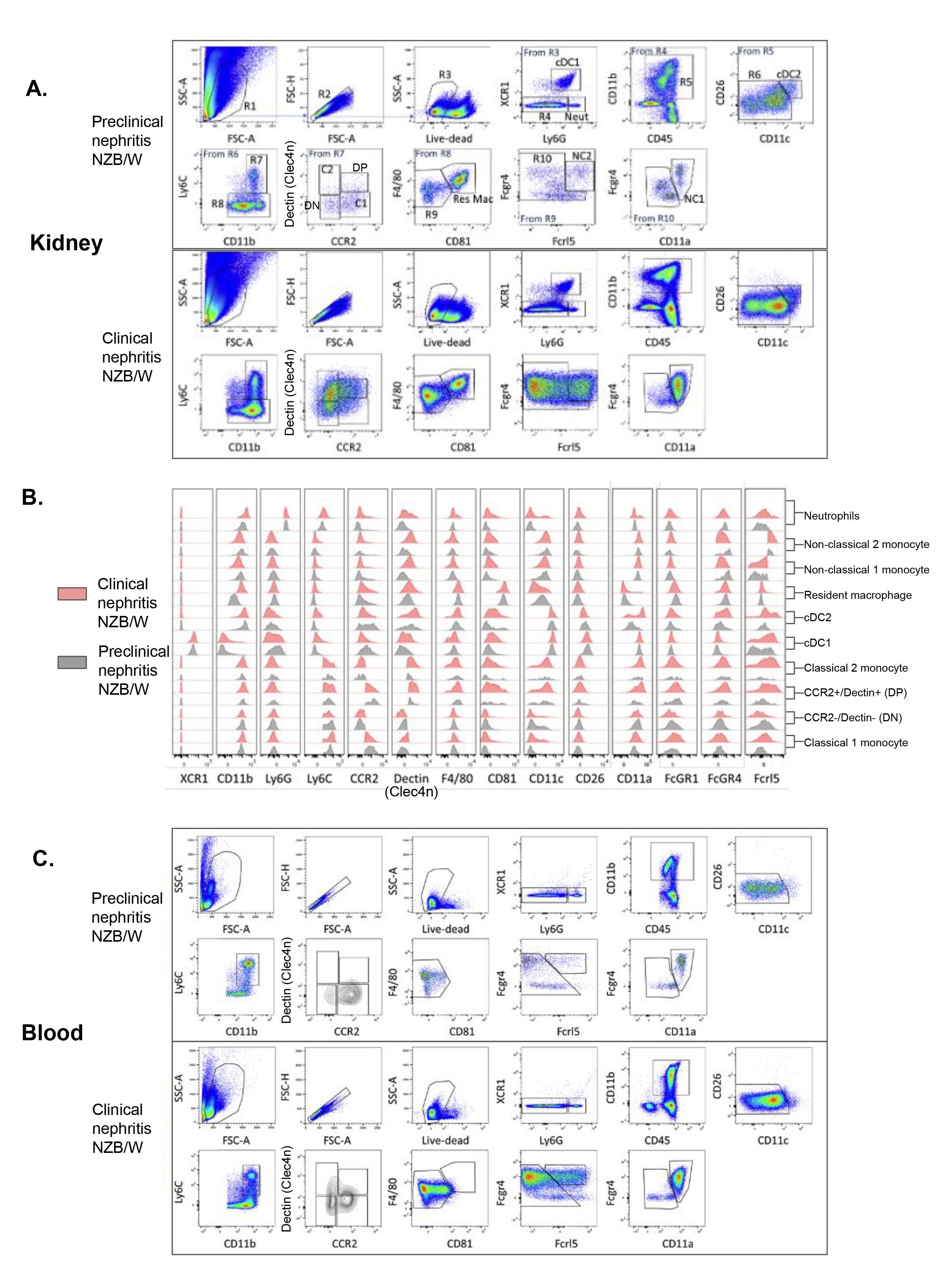
Gating strategy for sorting myeloid populations from mice that were identified by scRNA-seq in Figure 1. **A**. Sorting strategy for intrarenal myeloid populations. Using suspensions of kidney cells collected from mice with preclinical (top) or clinical (bottom) nephritis we positively selected cells using anti-CD45 beads. After gating for live single cells in the lymphocyte/myeloid gate, cDC1 and neutrophils were excluded using anti-XCR1 and anti-LyG respectively. The remaining cells were analyzed in the CD45+/CD11b+ gate (R5). cDC2 were identified as CD11chi/CD26+. Ly6Chi and Ly6Cint cells (R7) were subsetted using antibodies to Dectin2 and anti-CCR2. Classical 1 monocytes were defined as Ly6Chi/CCR2hi/Dectinlo and Classical 2 monocytes were defined as Ly6Cint/Dectin+/CCR2lo. Populations that were Dectin2lo/CCR2lo (DN) and Dectinhi/CCR2hi (DP) were also distinguished in the R7 gate. Ly6Clo cells (R8) were then used to gate resident macrophages (F4/80hi/CD81hi). Non-classical 2 cells were defined as Itgalhi/Fcgr4hi/Fcrl5hi and non-classical 1 cells were defined as Itgalhi/Fcgr4hi/Fcrl5lo. A small population of unidentified cells remained that comprised <5% of all myeloid cells. **B**. Expression histograms of antibody surface markers (x-axis) from intrarenal myeloid subsets shown from A (above). **C**. Sorting peripheral myeloid populations collected from mice with preclinical (top) or clinical (bottom) nephritis. We positively selected cells using anti-CD45 beads, gated on live single cells, and used a similar gating scheme as in A.

**Sup. Fig. 3.**
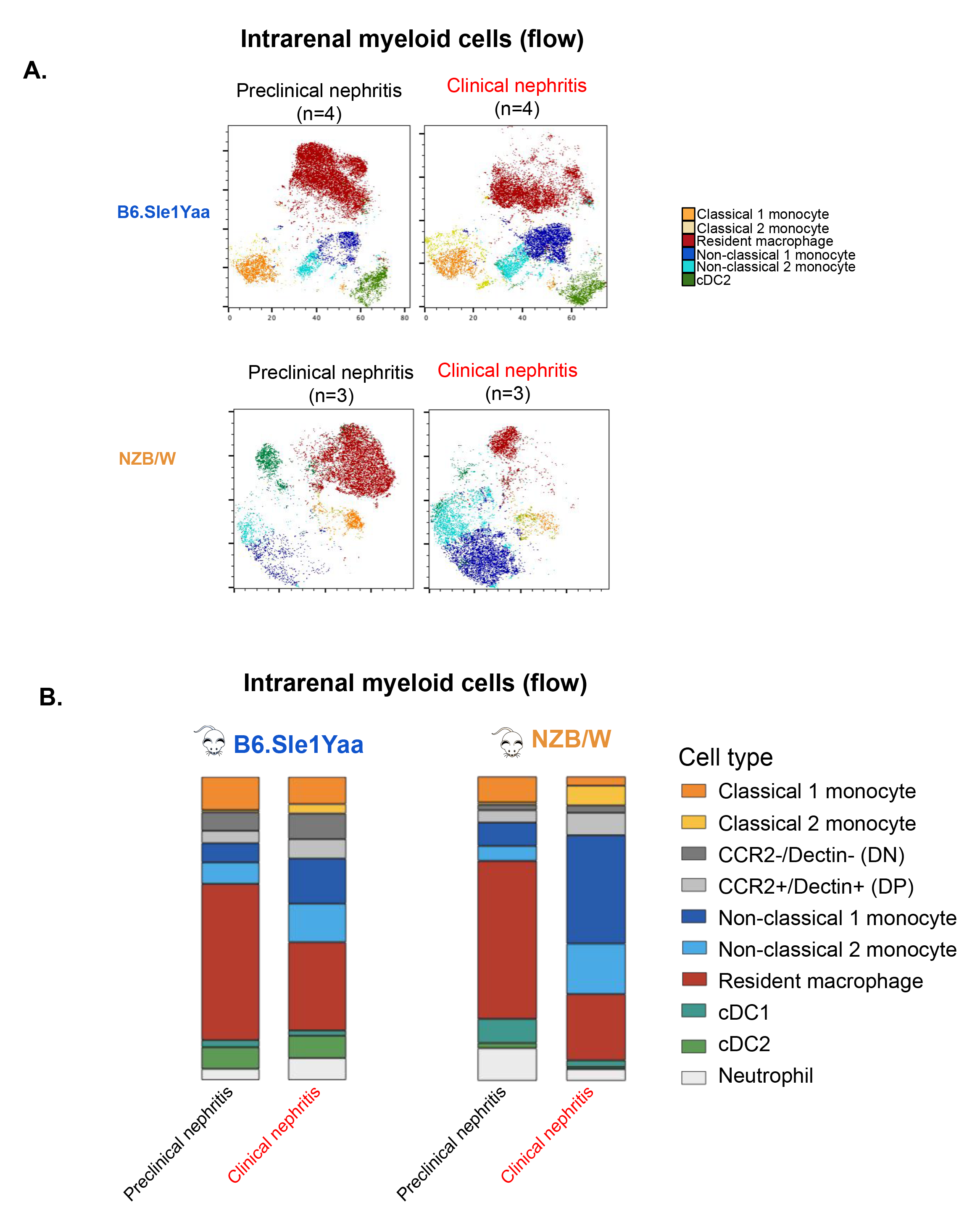
Monitoring intrarenal myeloid subsets from lupus mouse models during preclinical or clinical nephritis by flow cytometry. **A**. tSNE plots of intrarenal myeloid subsets in preclinical or clinical nephritis in NZB/W using flow cytometry. Each tSNE contains 16,500 cells from the CD45+ CD11b+ XCR1- Ly6G- gate (5,500 cells from each mouse). **B**. Average frequency of each myeloid subset measured via flow cytometry in B6.Sle1Yaa or NZB/W during preclinical or clinical nephritis. 5-8 mice were used for each condition.

**Sup. Fig. 4.**
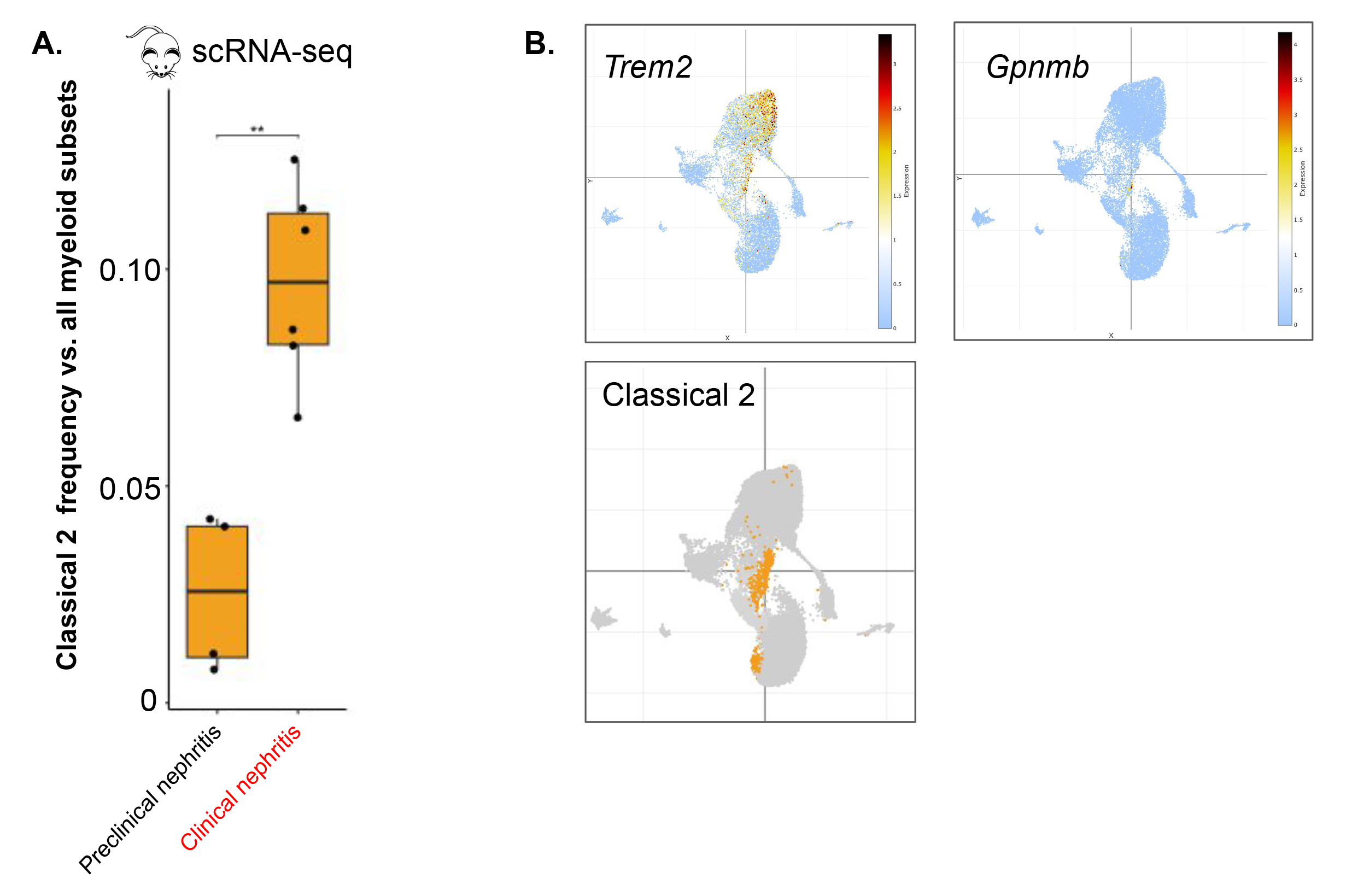
Classical 2 monocytes in mouse clinical nephritis. **A**. The frequency of Classical 2 monocytes relative to other myeloid cells in preclinical (n=4) or clinical nephritis (n=5) is determined by scRNA-seq droplet proportions. Each point represents C2 cells from 2-4 mice from the same strain that were experimentally pooled. **B**. Expression of injury associated genes in Classical 2 that were not differentially expressed relative to other myeloid clusters.

**Sup. Fig. 5.**
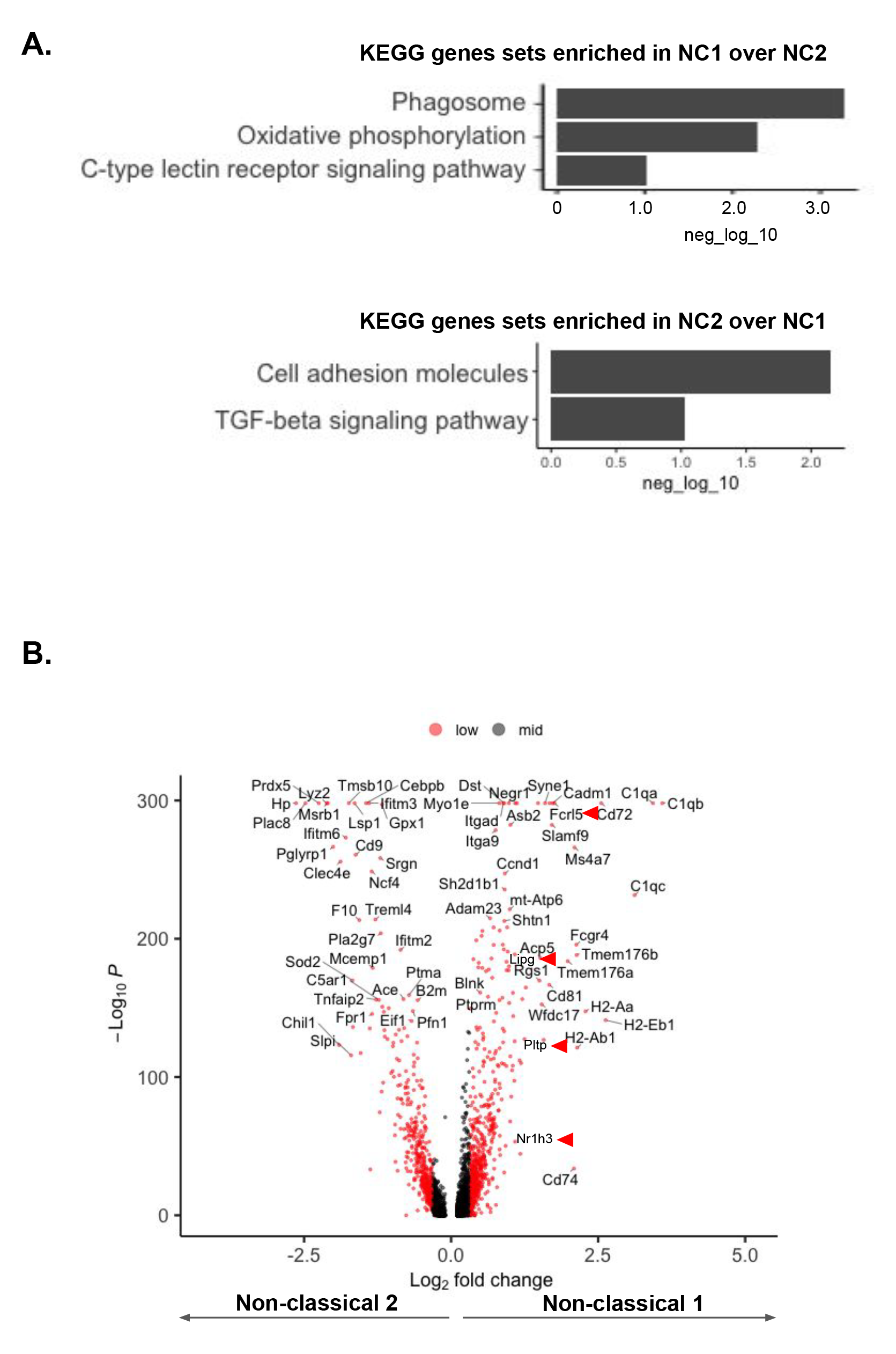
Non-classical 1 vs. non-classical 2 gene expression in mice. **A**. KEGG gene sets enriched in NC1 (top) or NC2 (bottom) from mice. **B**. Volcano plot depicting differentially expressed genes between Non-classical 1 and Non-classical 2 monocytes.

**Sup. Fig. 6.**
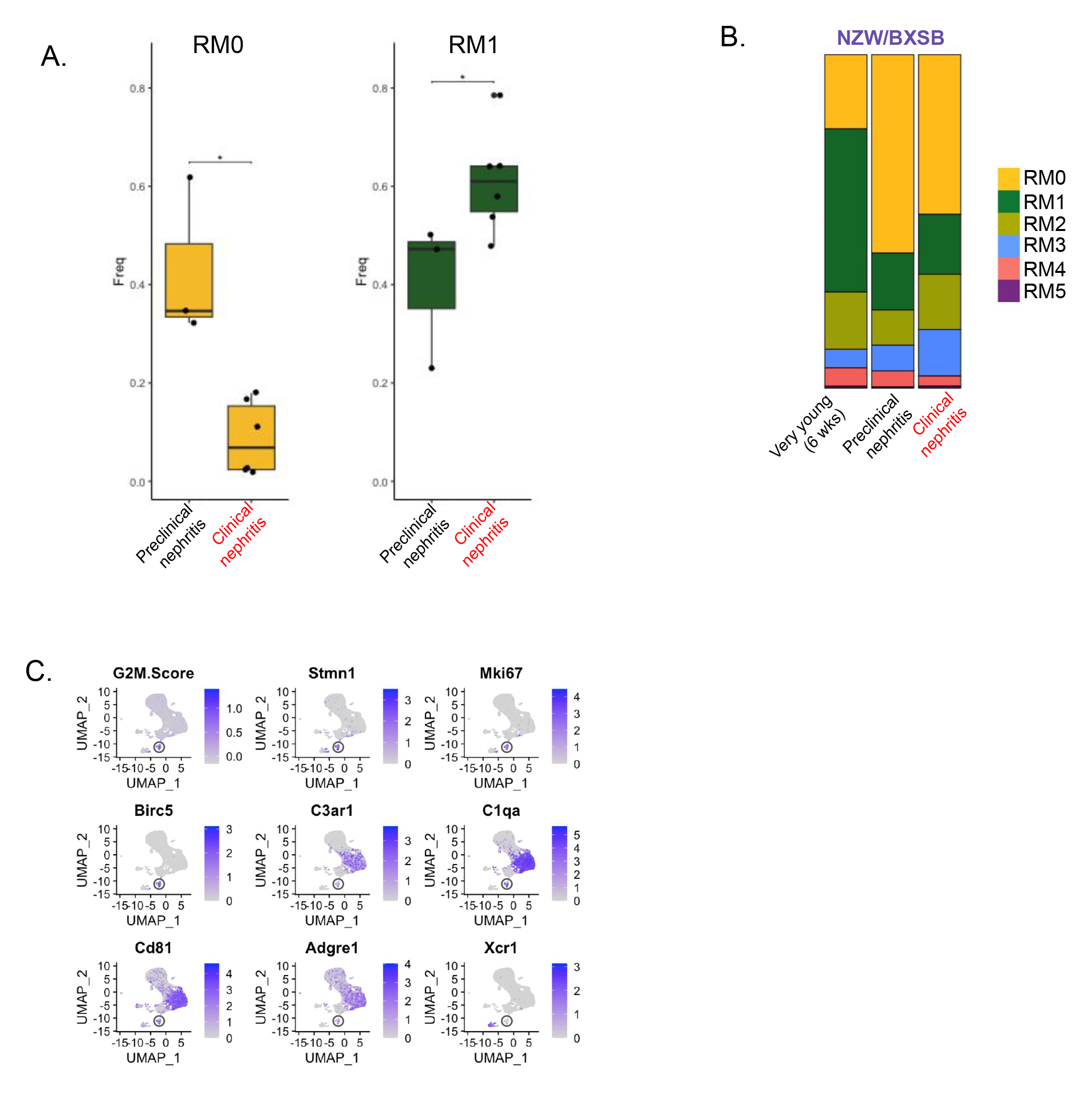
Summary of resident macrophage subsets from mice. **A**. The frequency of RM0 (left) and RM (right) in preclinical (n=4) or clinical nephritis (n=5) is determined by scRNA-seq droplet proportions. Each point represents RM cells from 2-4 mice from the same strain that were experimentally pooled. **B**. Normalized bar plots comparing myeloid subset frequencies in preclinical or clinical nephritis for NZW/BXSB at different time points as determined by scRNA-seq droplet proportions. **C**. Feature plots showing a cellular proliferation score (“G2M” using the Seurat cell cycle scoring function) and expression of canonical proliferation markers overlap. These overlap with a subset of resident macrophages (circled) identified by expression by expression of *C3ar1, C1qa, Cd81, Adgre1*. A small subset of DC1 shows a high G2M score.

**Sup. Fig. 7.**
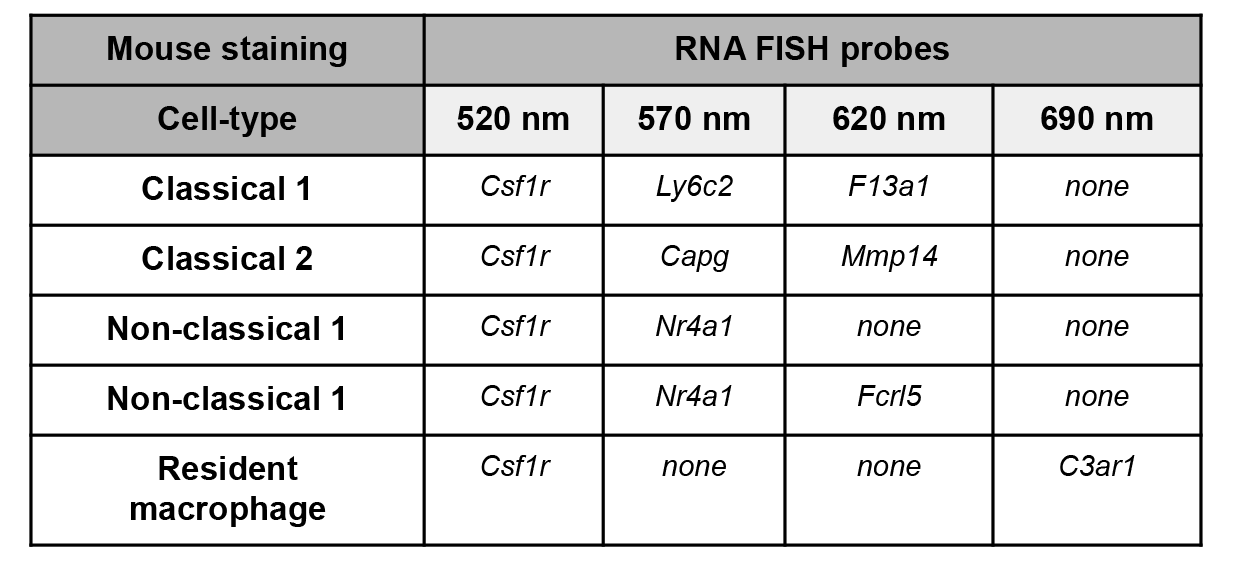
Table of FISH probes and wavelength used for staining and imaging the indicated cell type in mouse kidney sections.

**Sup. Fig. 8.**
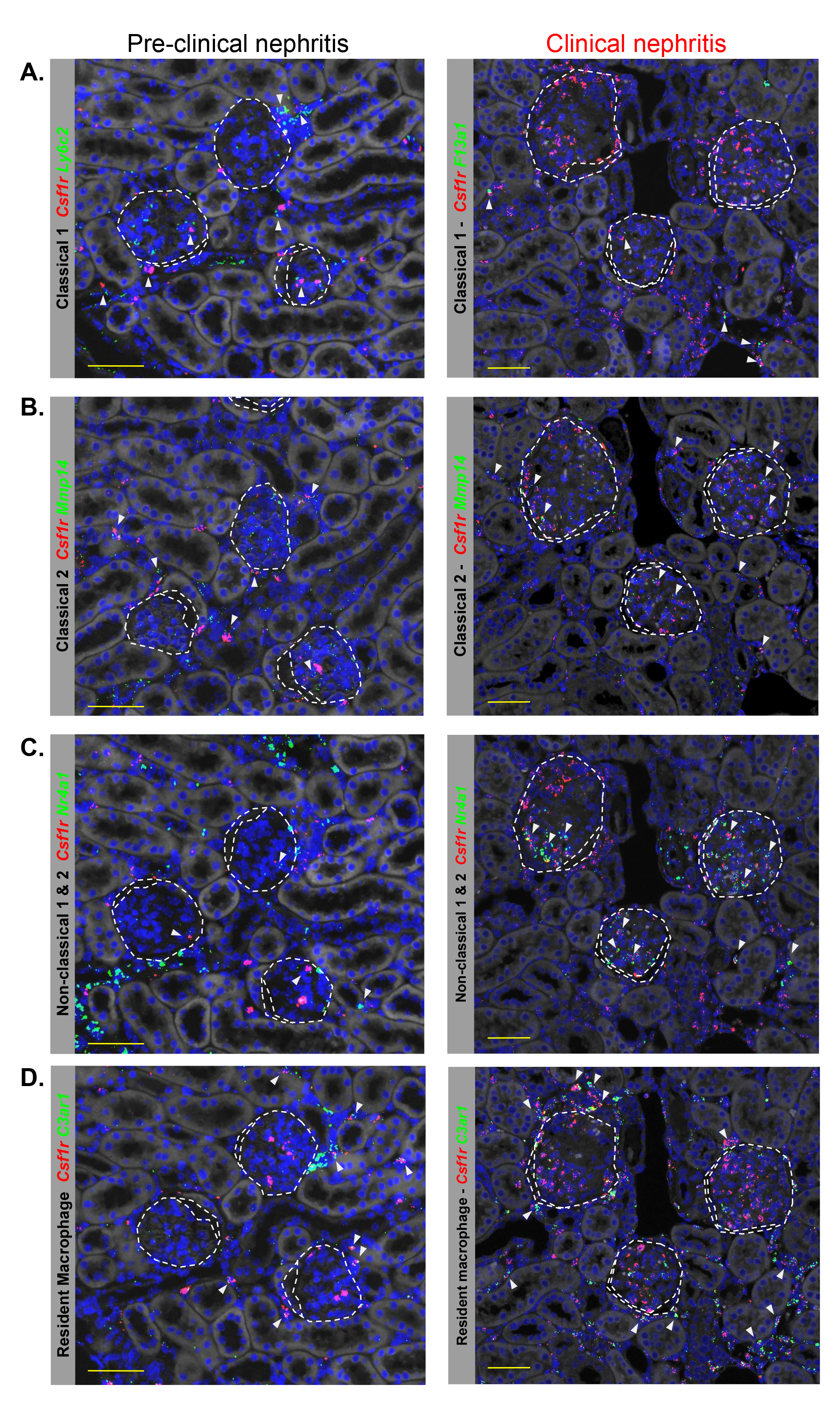
In situ localization of myeloid states identified by scRNA-seq in the cortex of serial kidney sections from Sle1.Yaa lupus mouse model. Kidney sections collected before (left) and after (right) clinical disease were stained with the indicated RNA FISH probes to identify (A) Classical 1, (B) Classical 2, (C) Non-classical, (D) Resident Macrophages. Arrows mark examples of the indicated cell type. Dotted lines represent outlines of glomerular tuft or adjacent urinary space within Bowman’s capsule. Scale bars represent 50 microns. We did not distinguish between intraglomerular Non-classical 1 vs 2 monocytes. Arrows mark examples of the indicated cell type. Scale bars represent 50 microns.

**Sup. Fig. 9.**
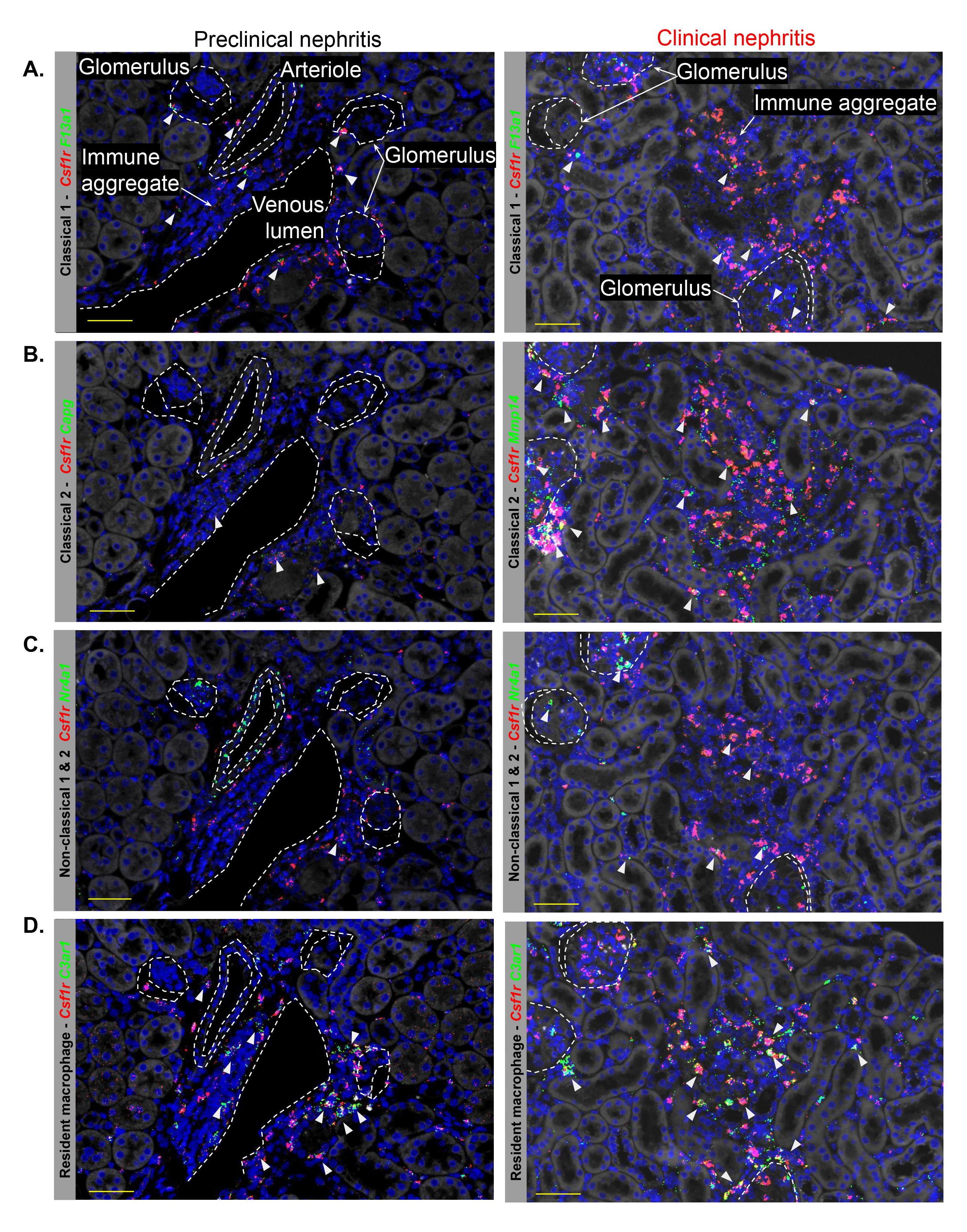
In situ localization of extra-glomerular myeloid states identified by scRNA-seq in serial kidney sections from Sle1.Yaa lupus mouse model. Kidney sections collected before (left) and after (right) clinical disease were stained with the indicated RNA FISH probes to identify (A) Classical 1, (B) Classical 2, (C) Non-classical, (D) Resident Macrophages. Note we did not distinguish between intraglomerular Non-classical 1 vs 2 monocytes. Dotted lines represent outlines of glomerular tuft or adjacent urinary space within Bowman’s capsule. Scale bars represent 50 microns. We did not distinguish between intraglomerular Non-classical 1 vs 2 monocytes. Arrows mark examples of the indicated cell type. Scale bars represent 50 microns.

**Sup. Fig. 10.**
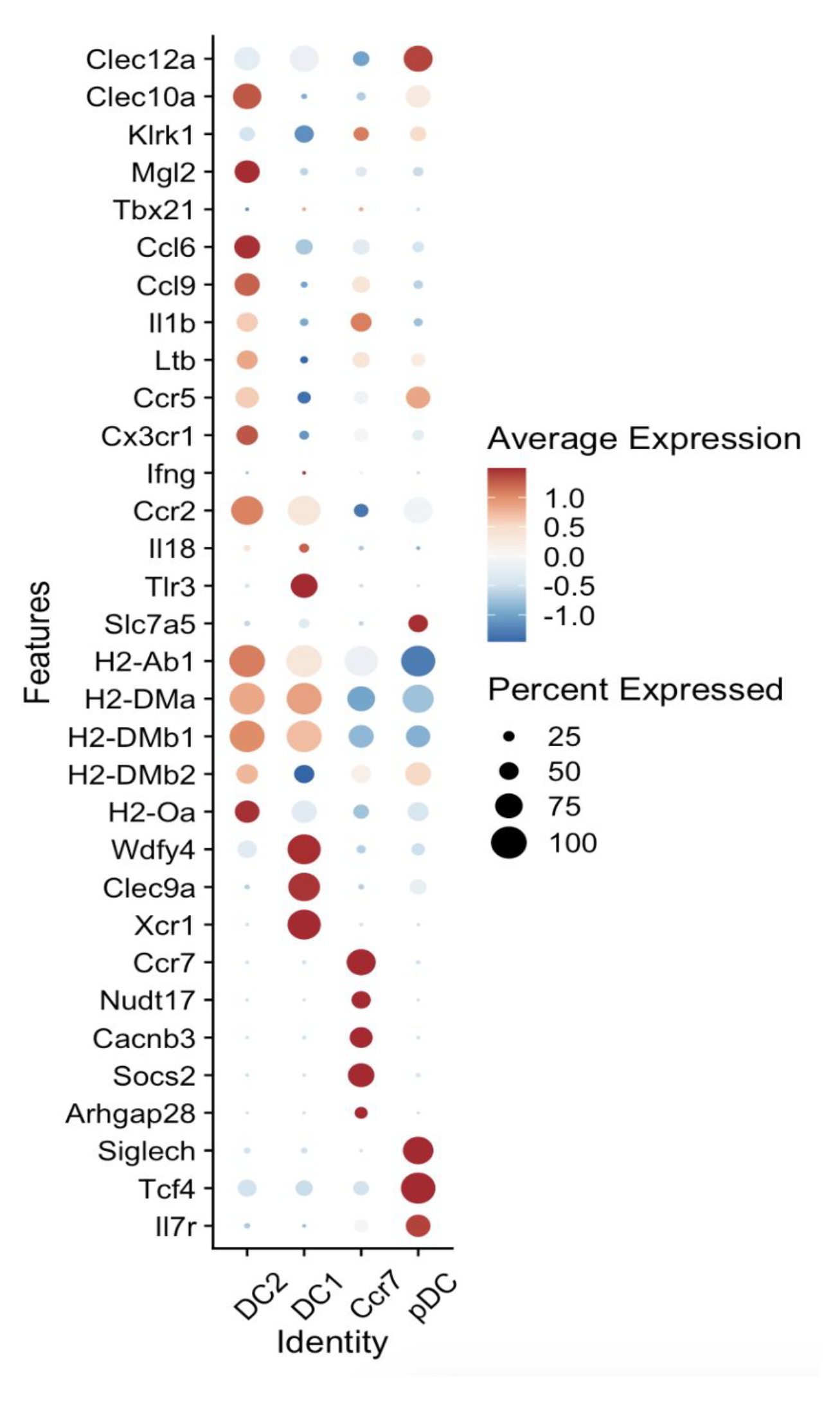
Expression of curated functional gene sets in dendritic cell subsets. Color scheme is based on z-score, from −1 (blue) to 1 (red).

**Sup. Fig. 11.**
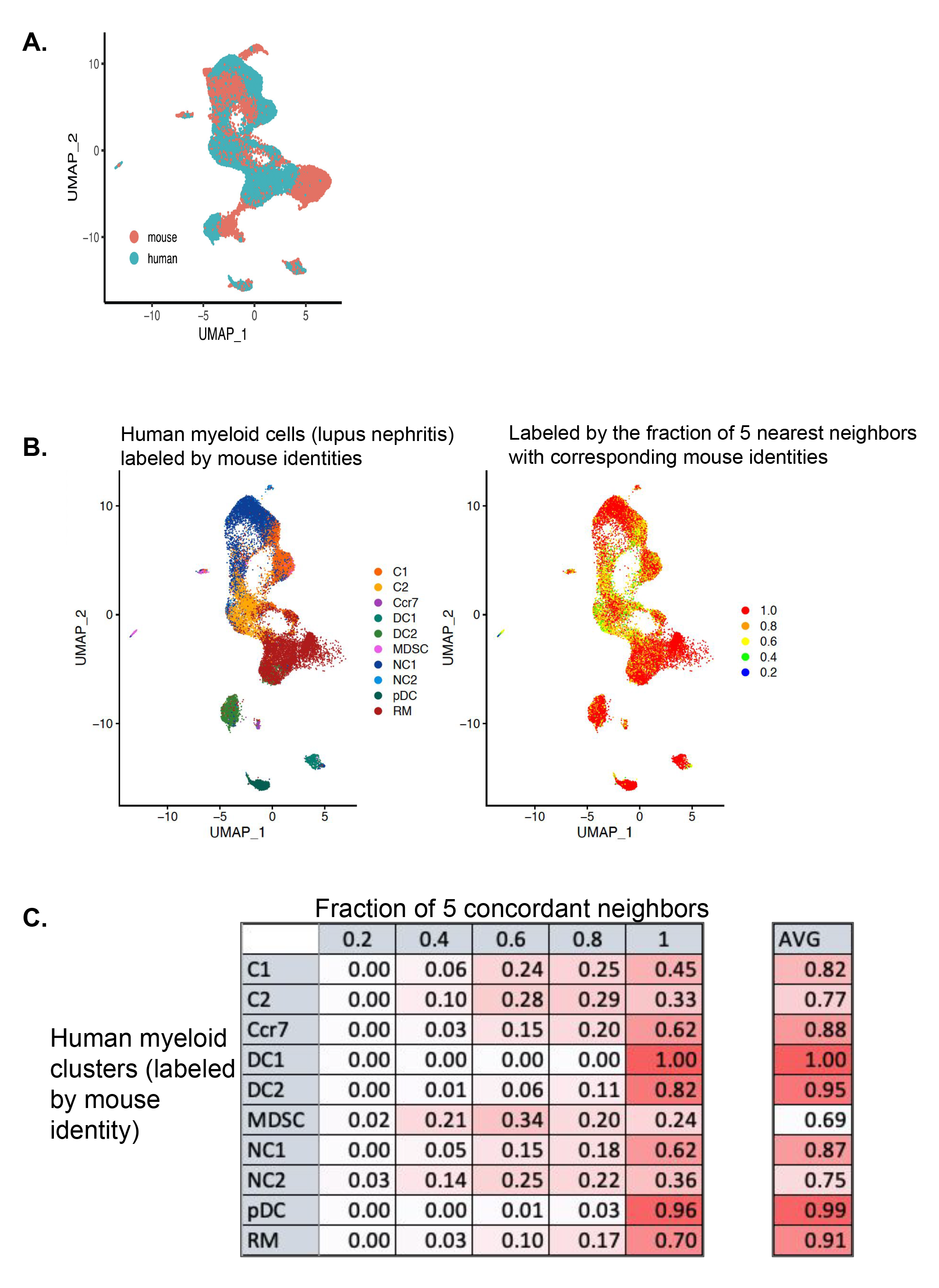
Integration of intrarenal myeloid sets from lupus mouse models and lupus patients. **A**. Harmonized UMAP embedding of myeloid cells from lupus mice and patients. **B**. Human cells were assigned with mouse identities (left) based on the correspondence of the 5 nearest neighbors. For each cell, we calculated the fraction of 5 nearest neighbors with corresponding mouse identities (right). **C**. Fraction of each human cluster with the indicated amount of concordant neighbors. The average concordance for all cells from each cluster is depicted on the right.

**Sup. Fig. 12.**
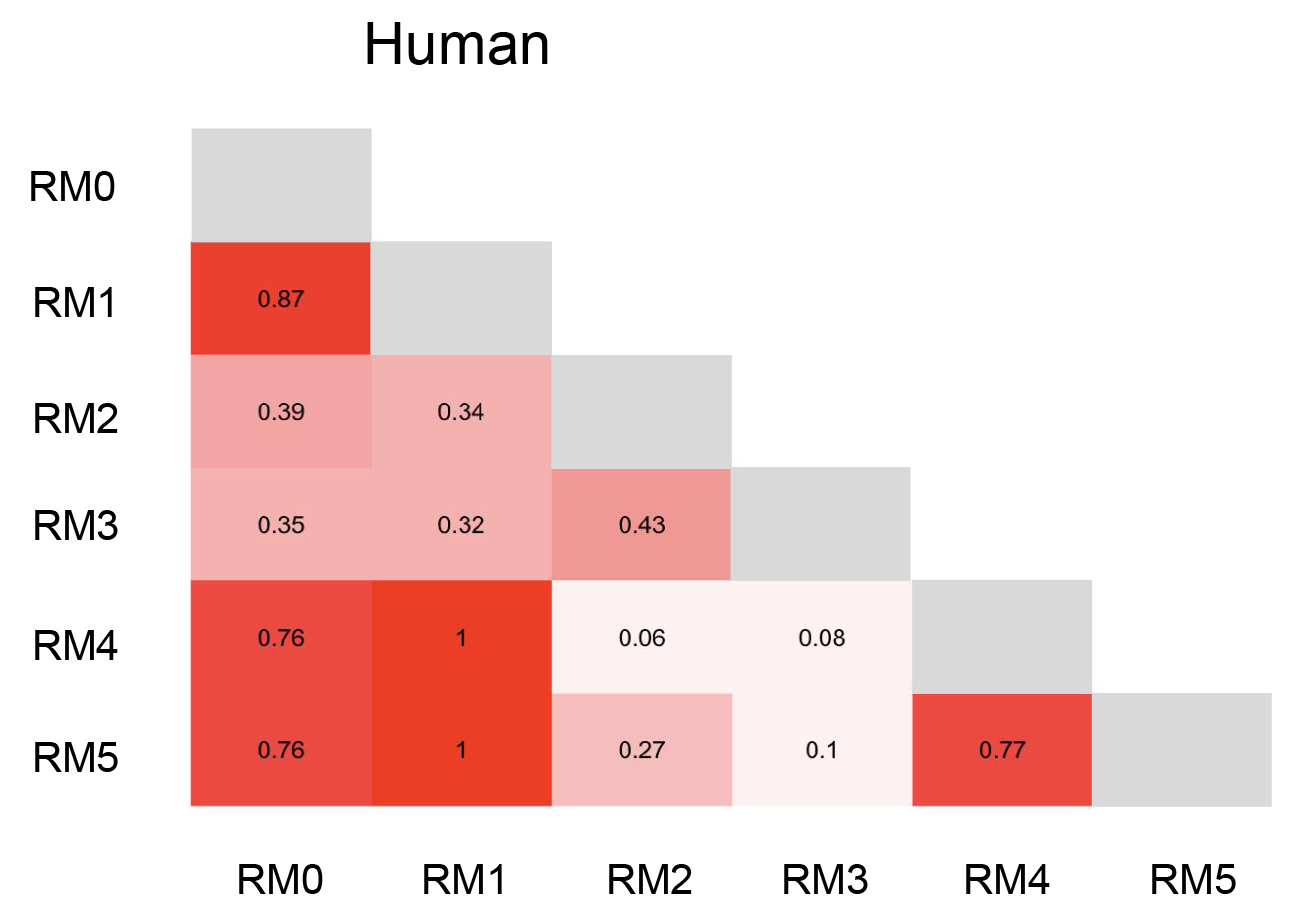
Summary of resident macrophage subsets from humans. **A**. Pairwise comparison of PAGA transcriptional relatedness of myeloid subclusters that were assigned RM mouse identities using KNN-label transfer. **B**. Scatter plots comparing the expression of marker genes from equivalent RM subclusters in mice (y-axis) vs. humans (x-axis). The right upper quadrant identifies conserved genes in the indicated RM subcluster.

**Sup. Fig. 13.**
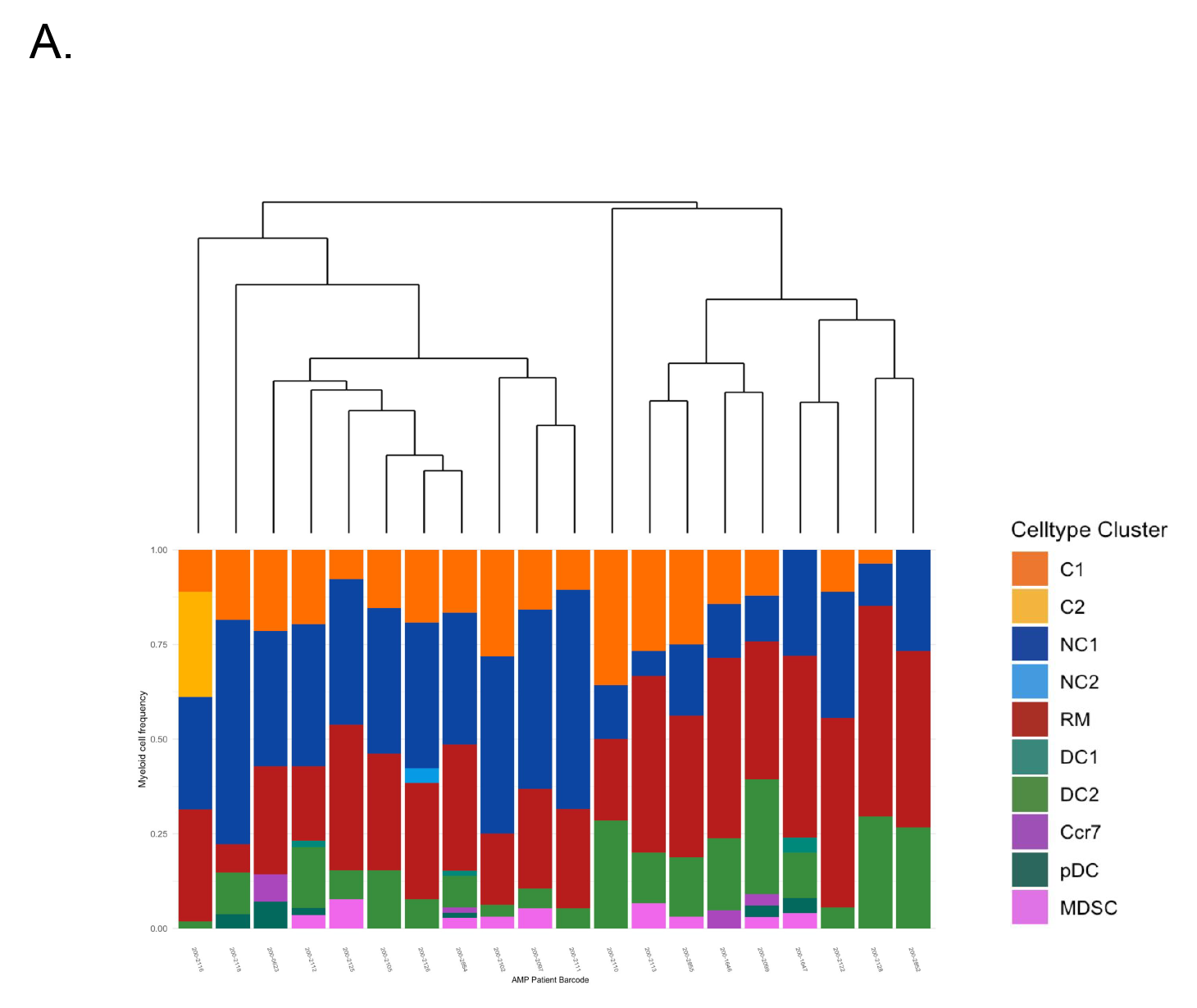
Intrarenal classical 2 monocytes are infrequent in healthy human kidneys. Normalized bar plots depicting the relative frequency of myeloid subsets in healthy control patients (n=20 patients with >10 myeloid cells per sample).

**Sup. Fig. 14.**
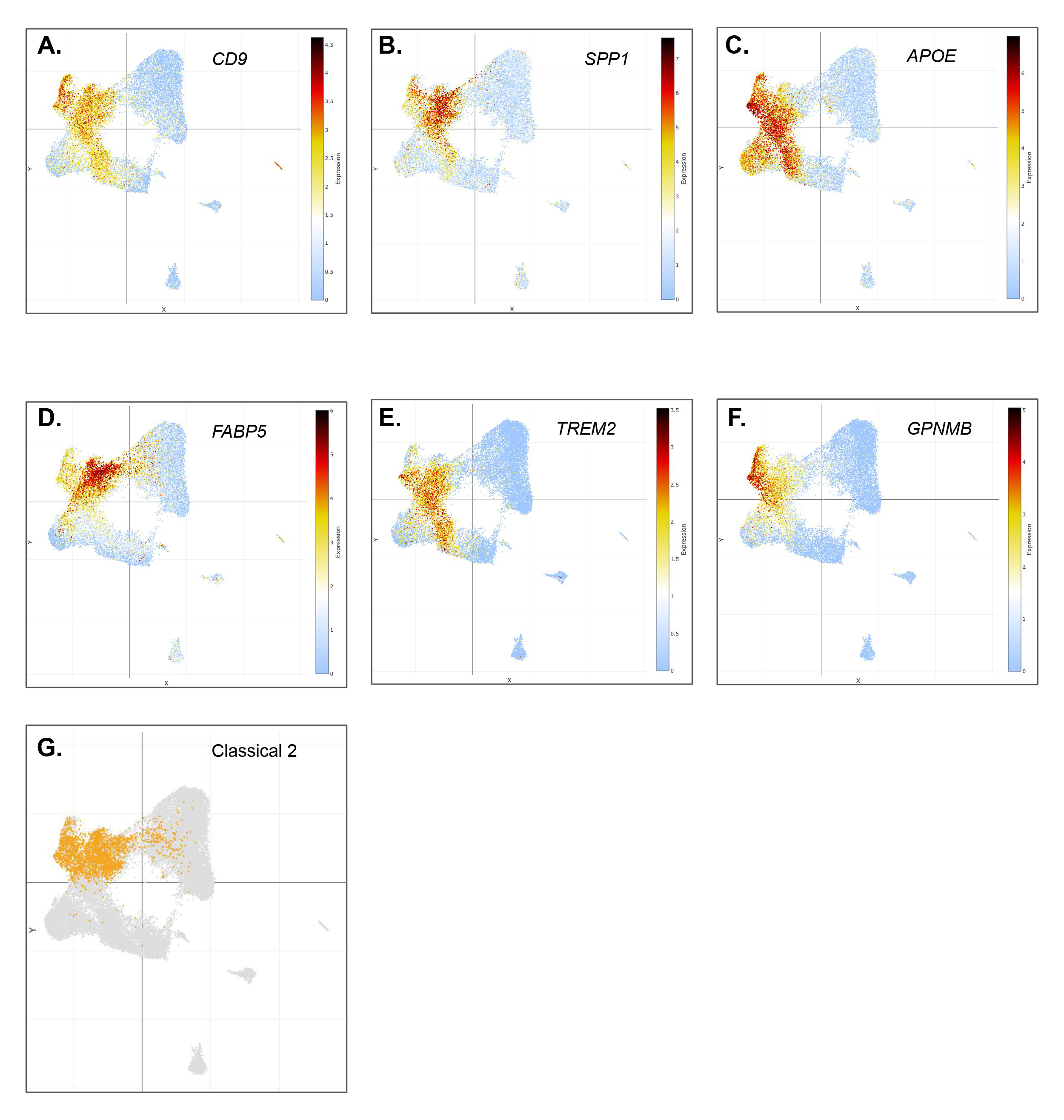
Expression of injury-associated genes in UMAP embedding of human intrarenal myeloid Cells. (A-F) (n=155 lupus patients). Overlay of classical 2 monocytes (G).

**Sup. Fig. 15.**
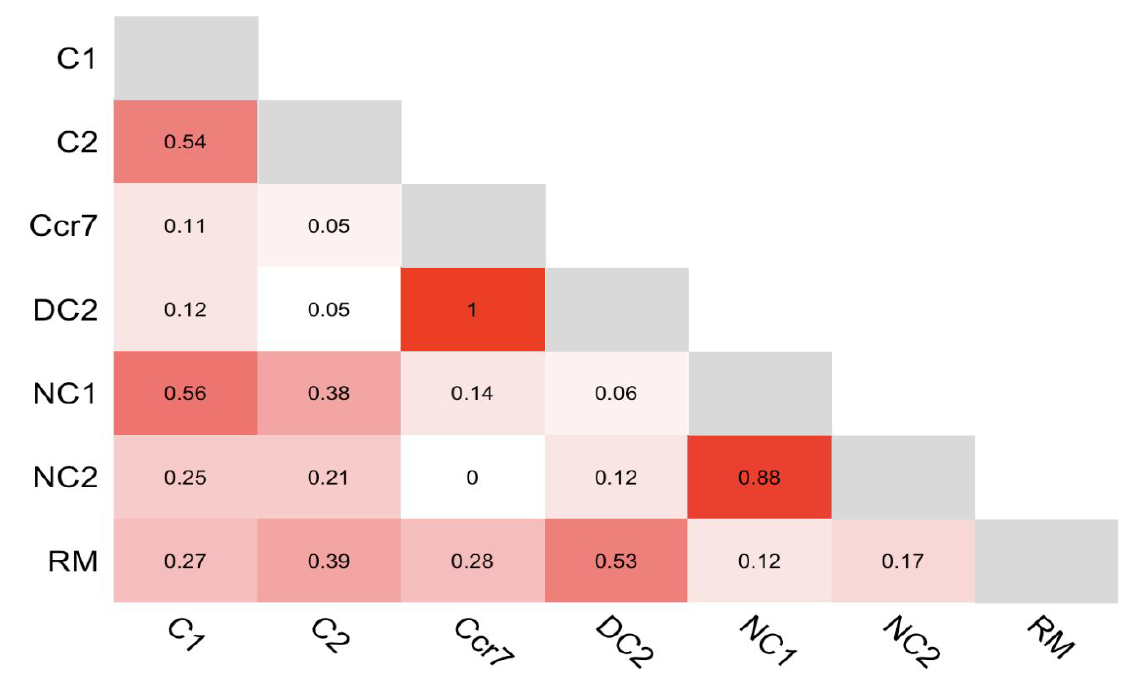
Pairwise comparison of PAGA transcriptional relatedness of human myeloid subclusters that were assigned mouse identities using KNN-label transfer.

**Sup. Fig. 16.**
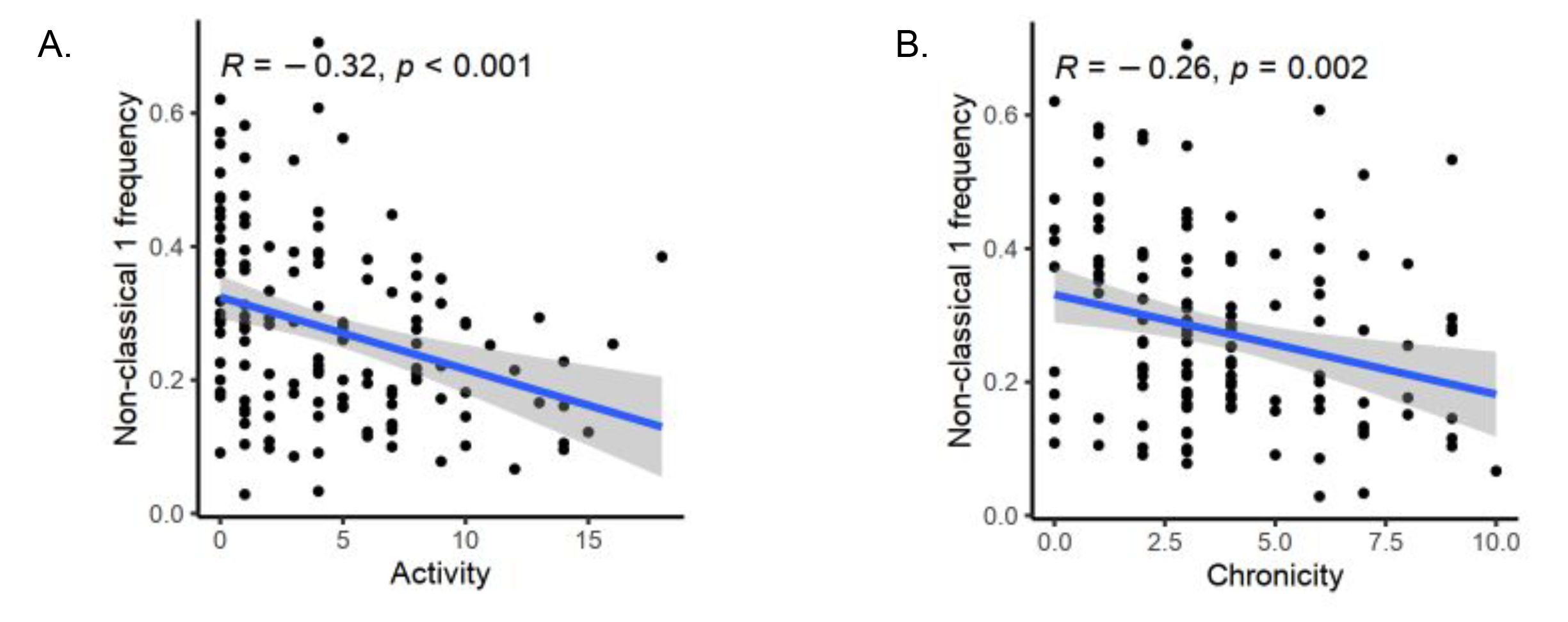
The frequency of human non-classical 1 is inversely associated with the activity (A) and chronicity (B) index.

**Sup. Fig. 17.**
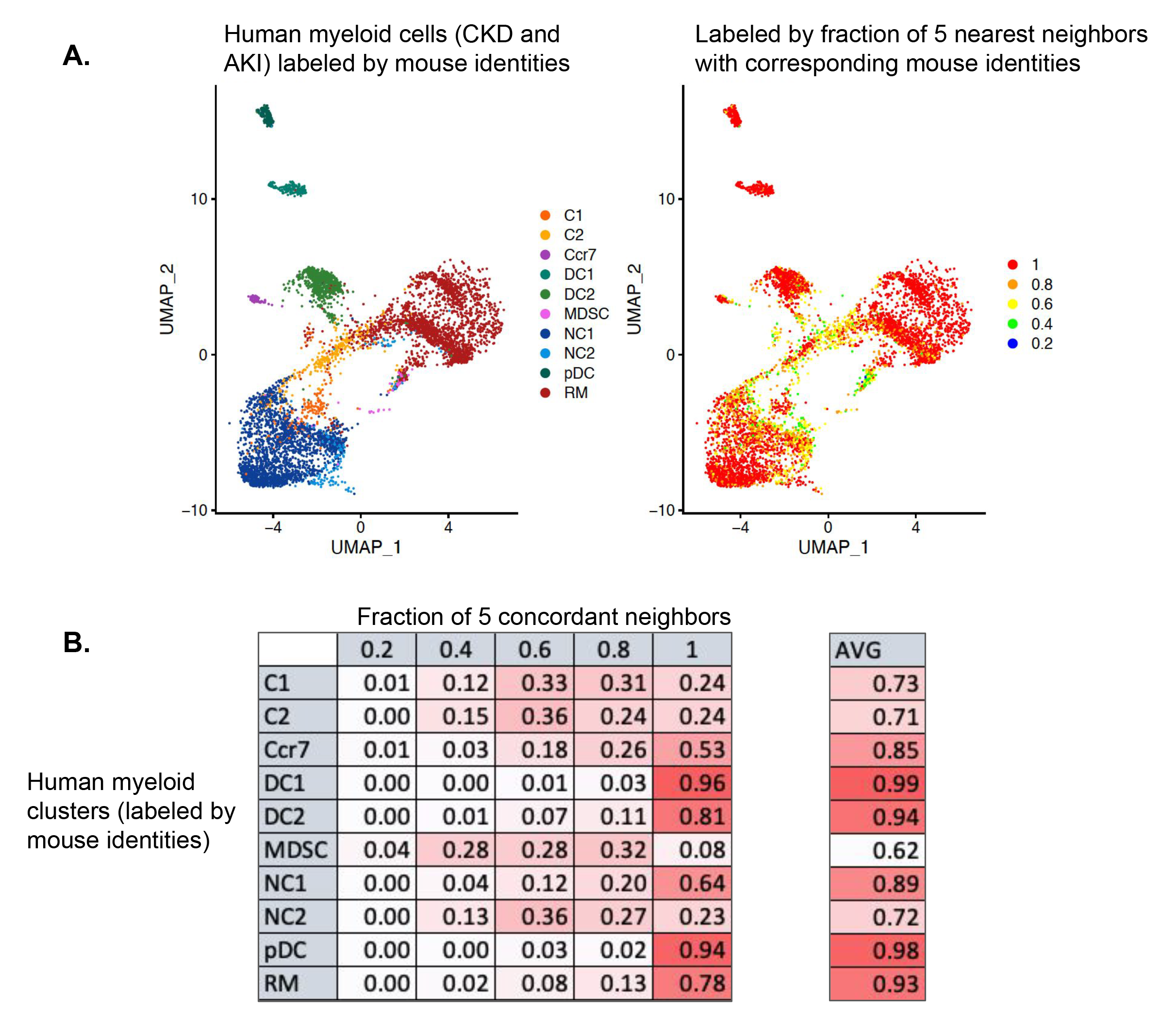
Integration of intrarenal myeloid sets from lupus mouse models and human patients with chronic and acute kidney injury. **A**. Human cells were assigned with mouse identities (left) based on the correspondence of the 5 nearest neighbors. For each cell, we calculated the fraction of 5 nearest neighbors with corresponding mouse identities (right). **B**. Fraction of each human cluster with the indicated amount of concordant neighbors. The average concordance for all cells from each cluster is depicted on the right.

**Sup Fig. 18.**
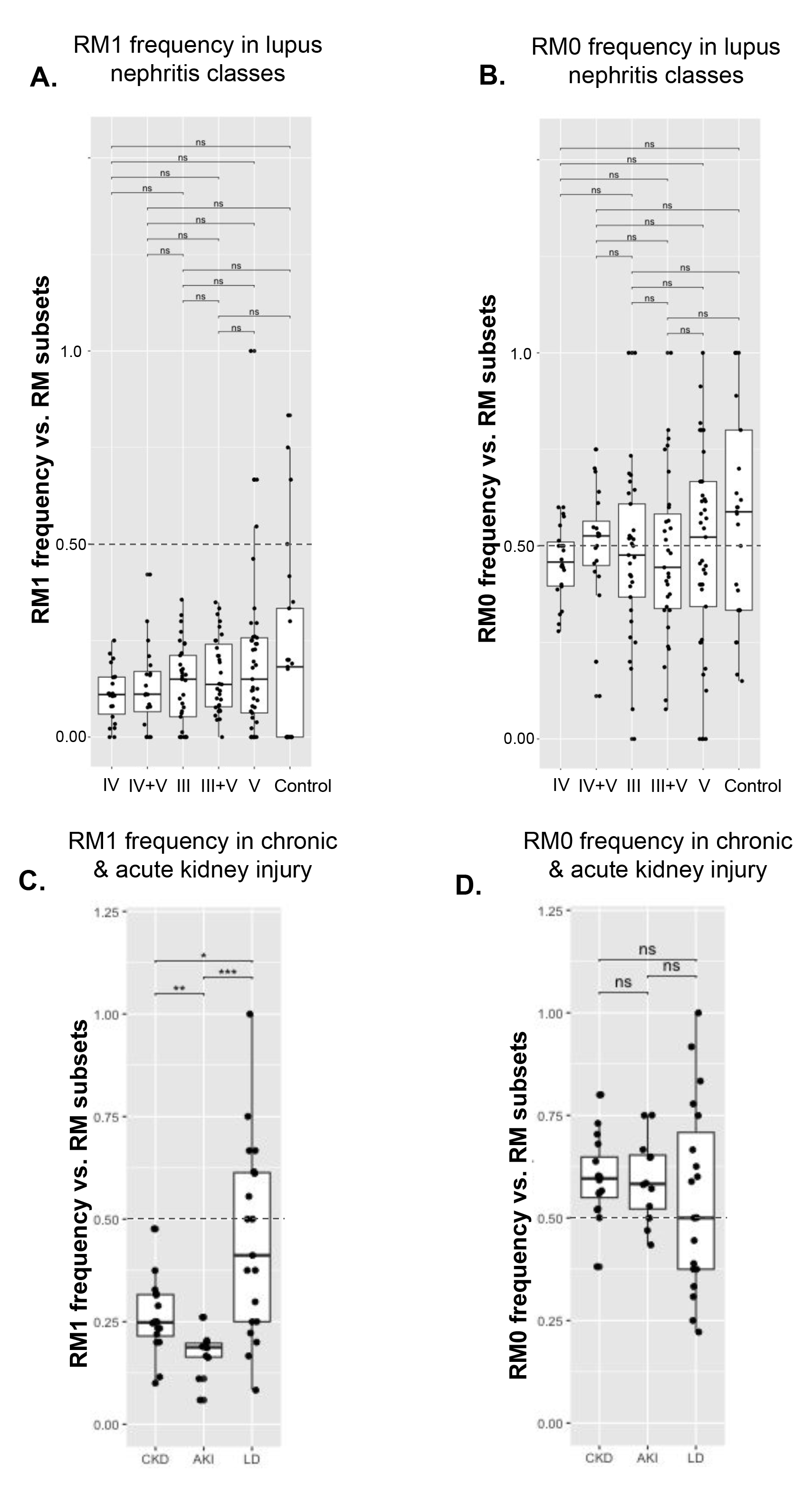
Human RM1 and RM0 frequencies relative to other RM subsets in patients with lupus nephritis. (A, B) and chronic an acute (C, D) kidney injury. Kruskal-Wallis test comparing means (ns: p > 0.05; *: p < 0.05; **: p < 0.01; ***: p < 0.001; ****: p < 0.0001).

**Sup Fig. 19.**
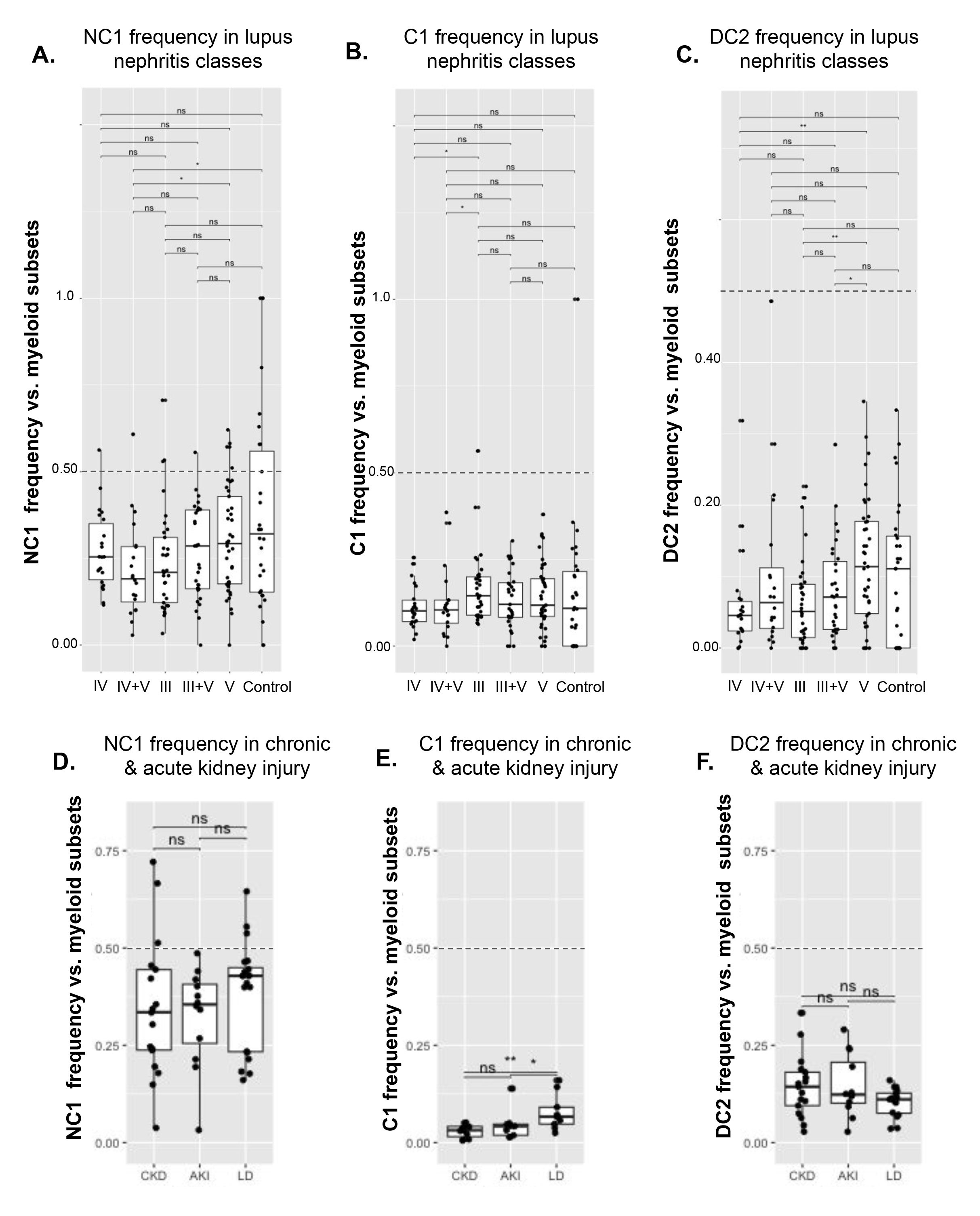
Human NC1, C1, and DC2 frequencies relative to other myeloid subsets in patients with lupus nephritis (A, B, C) and chronic an acute (D, E, F) kidney injury. Kruskal-Wallis test comparing means (ns: p > 0.05; *: p < 0.05; **: p < 0.01; ***: p < 0.001; ****: p < 0.0001).

**Sup. Fig. 20.**
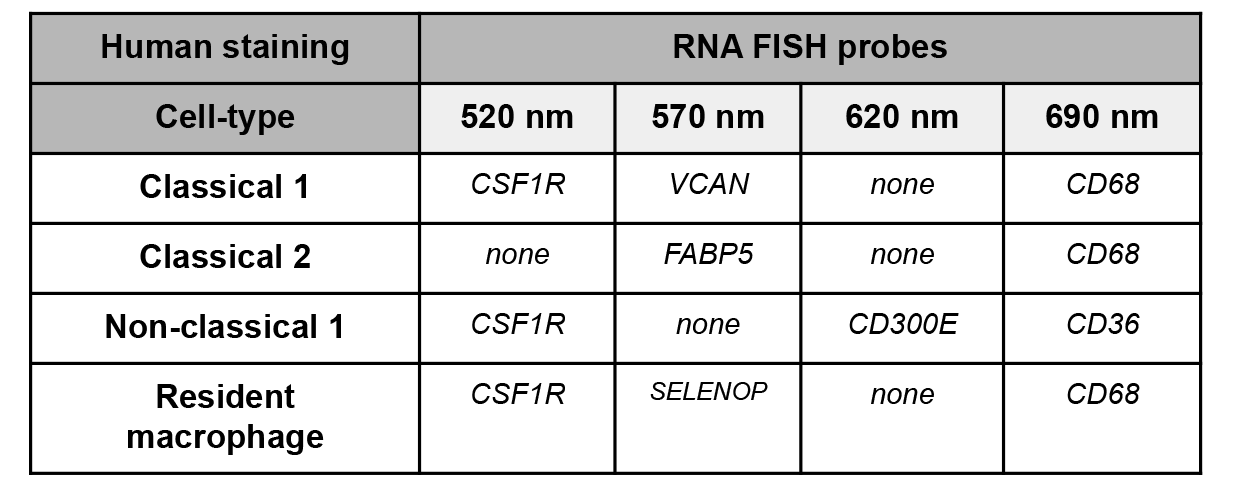
Table of FISH probes and wavelength used for staining and imaging the indicated cell type in human kidney sections.

**Sup. Fig. 21.**
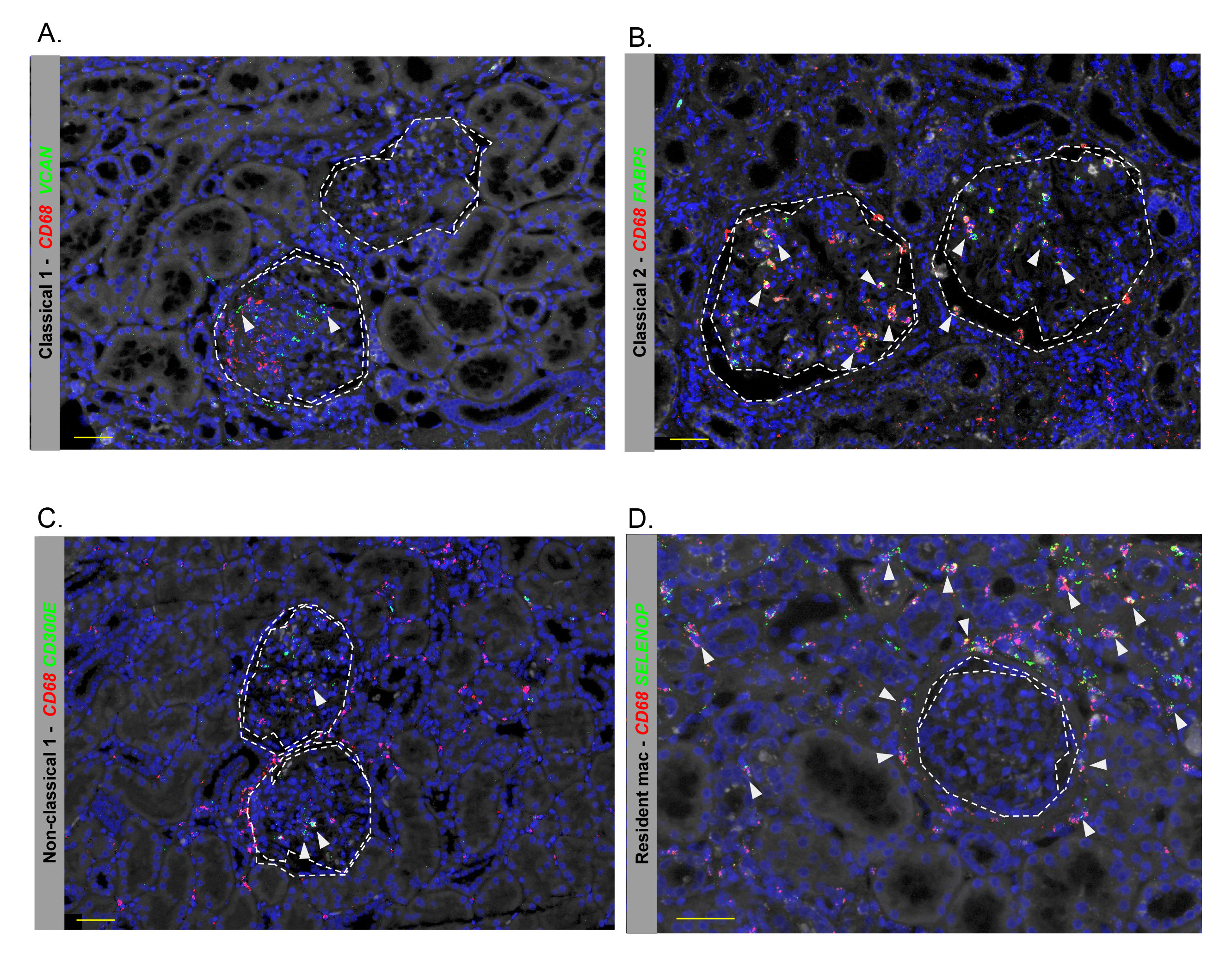
Representative *in situ* localization of myeloid states identified by scRNA-seq in glomerular and peri-glomerular areas of kidney sections from a human patient with lupus nephritis. Kidney sections were stained with the indicated RNA FISH probes to identify (A) Classical 1, (B) Classical 2, (C) Non-classical, (D) Resident Macrophages. Dotted lines represent outlines of glomerular tuft or adjacent urinary space within Bowman’s capsule. Scale bars represent 50 microns.

**Sup. Fig. 22.**
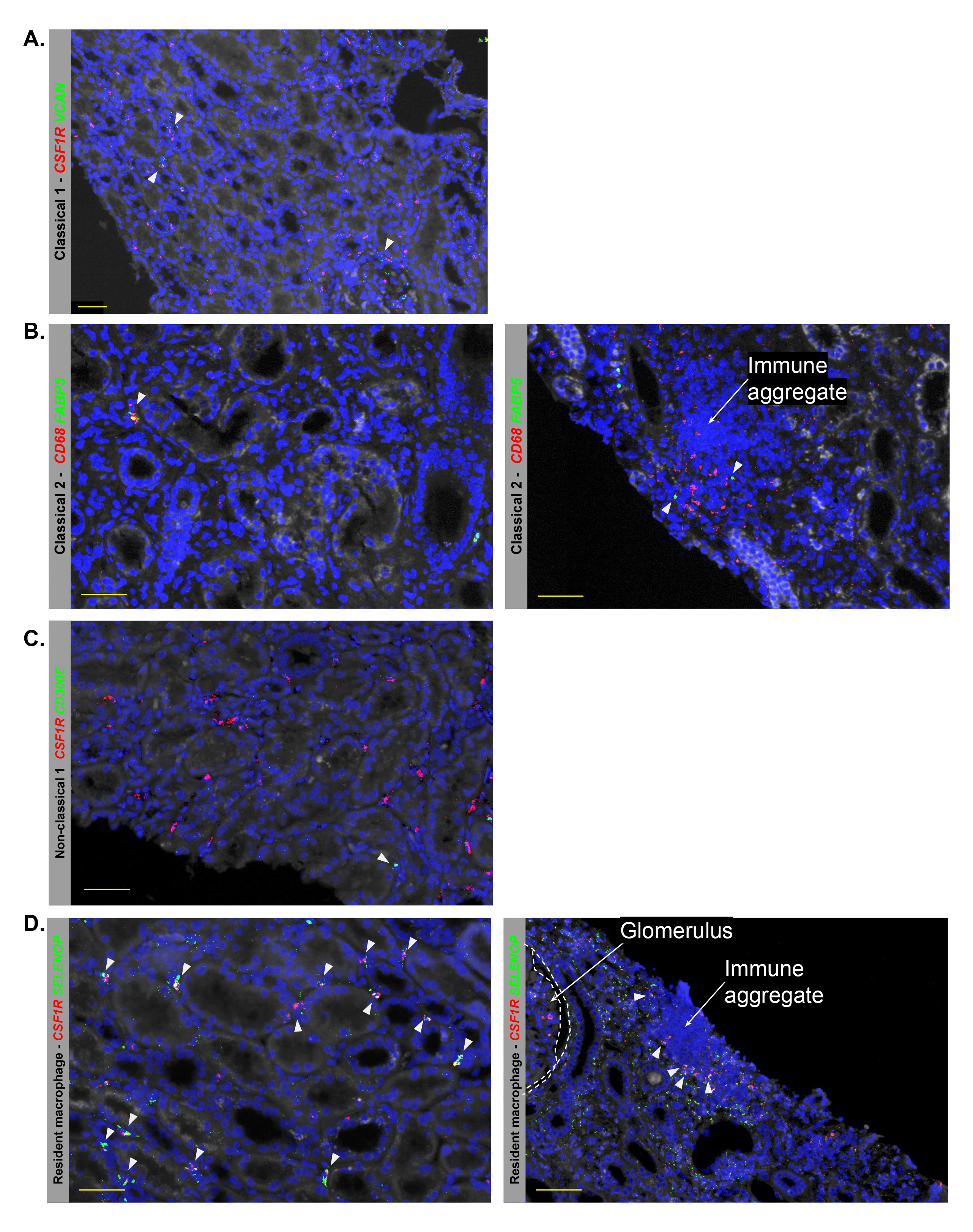
Representative *in situ* localization of myeloid states identified by scRNA-seq in extra-glomerular areas of kidney sections from a human patient with lupus nephritis. Kidney sections were stained with the indicated RNA FISH probes to identify (A) Classical 1, (B) Classical 2, (C) Non-classical, (D) Resident Macrophages. Dotted lines represent outlines of glomerular tuft or adjacent urinary space within Bowman’s capsule. Scale bars represent 50 microns.

## Notes

### Competing Interest Statement

The authors have declared no competing interest.

